# Alpha-toxin elicited CX_3_CL1-release via ADAM10 in *Staphylococcus aureus* pneumonia impairs bactericidal function of human monocytes

**DOI:** 10.1101/2023.09.17.554633

**Authors:** Srikanth Mairpady Shambat, Puran Chen, Rocky M. Barilla, Markus Huemer, Johanna Snäll, Amanda Welin, Taylor S. Cohen, Virginia Takahashi, Samuel B. Berry, Alejandro Gómez-Mejia, Tiziano A. Schweizer, Danen M Cunoosamy, Sara Cajander, Magda Lourda, Volkan Özenci, Ewerton Marques Maggio, Reto A. Schüpbach, Kristoffer Strålin, Annelies S. Zinkernagel, Anna Norrby-Teglund, Mattias Svensson

## Abstract

*Staphylococcus aureus* is an important human pathogen causing severe invasive infections. Pathogenesis is attributed to a wide array of virulence factors, including several potent exotoxins such as the pore-forming alpha-toxin. In this study, we found that patients with *S. aureus* respiratory tract infections had elevated CX_3_CL1 levels in airway fluid and plasma. Using humanized organotypic lung models, we observed that stimulation of lung epithelium with alpha-toxin induce an intensified CX_3_CL1 expression apically in the epithelium as well as the release of CX_3_CL1. Blocking alpha-toxin or ADAM10 activity in organotypic lung using an alpha-toxin-blocking antibody or a specific ADAM-10 inhibitor confirmed their role in modulating CX_3_CL1 cleavage and release. Analyses of CD14^+^ human monocytes in combination with a CX_3_CR1 inhibitor revealed that alpha-toxin-mediated CX_3_CL1 release induce CX_3_CL1-dependent chemotaxis. In line with these data, lung tissue from patients with *S. aureus* respiratory tract infection showed elevated CX_3_CL1 and CD14 staining as compared to tissue from patients with non-infectious lung diseases. Functional studies of monocytes showed that CX_3_CL1 released by lung models resulted in upregulated CD83 and downregulated CD86, as well as impaired killing of phagocytosed *S. aureus*. Furthermore, stimulation of monocytes with soluble CX_3_CL1 hampered their reactive-oxygen and nitric-oxide production. Taken together, our data show that *S. aureus* triggers the release of lung epithelial CX_3_CL1; a process found to be dependent on the alpha-toxin’s effect on ADAM10 mediating cytotoxicity and resulting in impaired monocyte phagocytic killing. Hence, we identify an immunomodulatory effect of alpha-toxin involving the CX_3_CL1-ADAM10 axis extending beyond the cytolysis function.

## INTRODUCTION

The pathophysiology of *Staphylococcus aureus* pneumonia involves a variety of virulence mechanisms mediated by secreted degradative enzymes, immune evasion, and cytotoxic factors such as exotoxins (1–3). The pore-forming exotoxin α-toxin is one of the most well-studied virulence factors of *S. aureus*. It has been shown that α-toxin mediates cell damage directly via its pore-forming capability and contributes to lethality in pulmonary infection (4, 5). *S. aureus* strains lacking α-toxin show reduced cytotoxicity and virulence in invasive lung disease models and the more α-toxin *S. aureus* produces, the greater the airway epithelial damage (5, 6). Consequently, using mouse and rabbit *S. aureus* pneumonia model, α-toxin has been shown to be a key virulence determinant causing severe lung tissue pathology (7, 8). Following the identification of A Disintegrin and metalloproteinase domain-containing protein 10 (ADAM10) as a receptor for α-toxin (4), major advances have been made in understanding the pathogenesis and host cell specificity of *S. aureus*. Binding of α-toxin to ADAM10 expressed by epithelial cells (9), activation of intracellular signaling pathways following pore-formation have also been shown to increase the enzymatic activity of ADAM10 (10). The role of ADAM10 in α-toxin-mediated lethality has been partly attributed to the cleavage of E-cadherin in lung tissue of mice infected with *S. aureus* (11). Furthermore, the targeting of ADAM10 by α-toxin can hijack its regular homeostatic function by increasing its enzymatic activity and causing cleavage of tight junction proteins (12–14). Similarly, α-toxin-mediated epithelial damage is not only limited to its receptor engagement. Oligomerization of α-toxin to form a heptameric β-barrel pore in the membrane (15), followed by pore formation and influx of ions such as Ca^2+^ can lead to cell lysis (9, 12, 14, 16). Subsequent pore-formation in the airway epithelial cells can lead to several downstream inflammatory responses such as increased levels of pro-inflammatory mediators, increased epithelial permeability and vascular leakage (17–19). Since there is an extensive list of potential ADAM10 substrates, perturbations to the tightly regulated activity of ADAM10 may have consequences in the pathogenesis of *S. aureus* pneumonia beyond ADAM10’s role as an α-toxin docking protein and its cleavage of E-cadherin (20). ADAM proteins have recently been reviewed and highlighted in the regulation of immune cells (21). One ADAM10 substrate that regulates immune cell function is the chemokine CX_3_CL1 (also known as fractalkine). CX_3_CL1 is produced by various cell types, including lung epithelial cells. In its membrane-bound form CX_3_CL1 promotes cell-cell adhesion and it induces chemotaxis as a soluble molecule. The proteolytic cleavage of membrane-bound CX_3_CL1 by a metalloproteinase is a unique mechanism of chemokine-release and has been implicated in recruitment and transmigration of immune cells to CX_3_CL1-expressing mucosal tissue such as the respiratory tract (22, 23). Increased levels of CX_3_CL1 have been reported in infectious and inflammatory diseases (24, 25) but whether *S. aureus* respiratory tract infections and α-toxin are associated with CX_3_CL1-release is not known. Additionally, whether α-toxin interaction with ADAM10 at sub-lytic concentrations in mucosal tissue modulate immune cell functions and contribute to the pathogenesis locally is poorly understood.

Here, we examined patients with *S. aureus* lung infections and used a human *in vitro* organotypic model of *S. aureus* pneumonia (6). We found that in *S. aureus* lung infection, release of lung epithelial membrane-bound CX_3_CL1 is dependent on α-toxin and ADAM10, and that soluble CX_3_CL1 modulates monocyte phenotype and function. This study reveals a α-toxin-ADAM10-CX_3_CL1-axis as a novel component of α-toxin-mediated modulation of effector functions of immune cells and of pathogenesis in *S. aureus* respiratory infections.

## RESULTS

### Local and systemic levels of CX_3_CL1 are increased in patients with *S. aureus* respiratory tract infection

To investigate the relationship between *S. aureus* respiratory tract infections and CX_3_CL1, plasma and lung airway fluids (bronchioalveolar lavage and tracheal aspirates) from prospectively enrolled patients with confirmed *S. aureus* respiratory tract infection were collected and analyzed for soluble CX_3_CL1. Compared to other severely ill non-infectious ICU prospectively enrolled patients or healthy volunteers, we observed on average 10 times higher levels of CX_3_CL1 in the *S. aureus* respiratory tract infections group in both lung fluids and plasma, respectively (Fig. 1 A and B, Supplementary Table 1 and Table 2). Next, we compared the levels of CX_3_CL1 in frozen plasma samples from healthy volunteers and a retrospective cohort of patients with confirmed *S. aureus* bloodstream infection, without respiratory tract infections. Healthy volunteers had undetectable levels of CX_3_CL1 (*p*=0.66, 14 out of 19 below limit of detection, LOD) while patients with *S. aureus* bloodstream infection had slightly elevated levels (*p*<0.01) (Fig. 1 C). Notably, compared to the patients with *S. aureus* respiratory tract infection, many of the patients with *S. aureus* bloodstream infection had undetectable levels of CX_3_CL1 (10 out of 23 below LOD). We observed no association between CX_3_CL1 levels and clinical parameters such as presence of co-morbidity, age or severity/outcome in the *S. aureus* patient cohort (Supplementary Fig. 1 A-C). Taken together, these data suggest that *S. aureus* respiratory tract infection increases CX_3_CL1 levels both locally and systemically.

**Figure 1.**
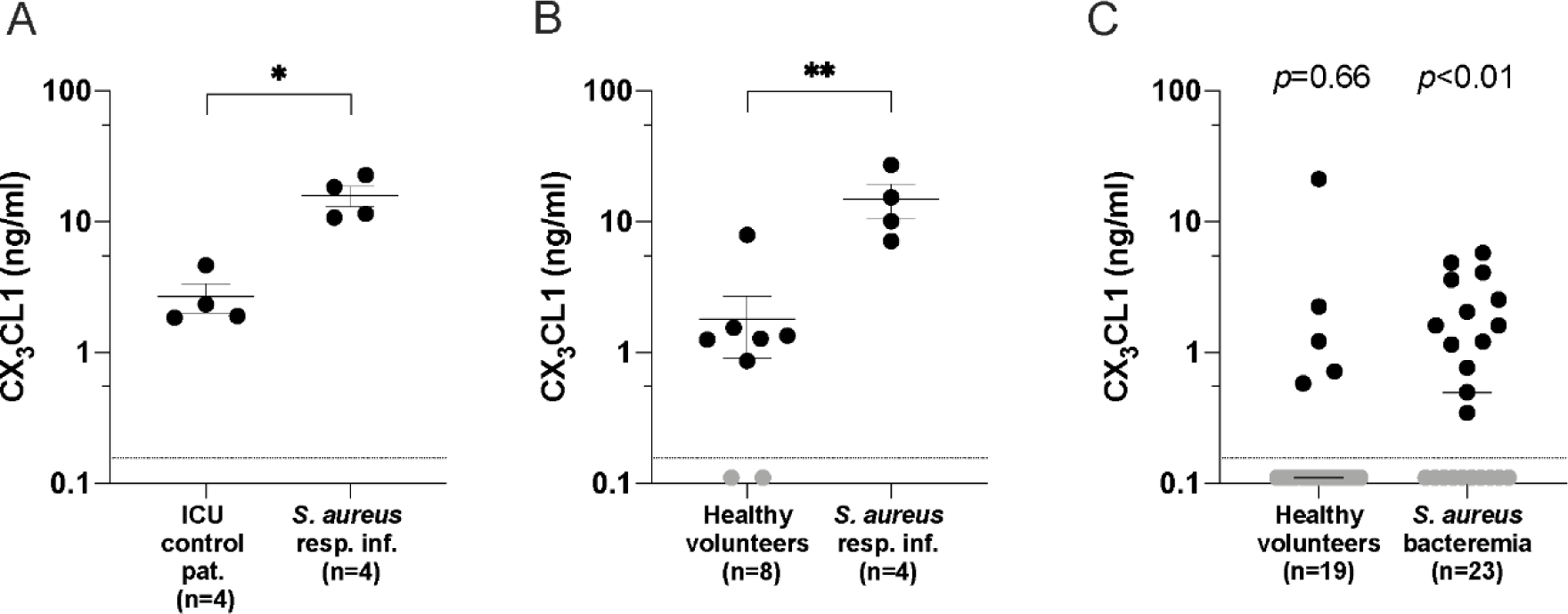
CX_3_CL1 levels in samples from *Staphylococcus aureus* infected patient. A-B) CX_3_CL1 protein levels in lung airway fluids (bronchioalveolar lavage and tracheal aspirates) from ICU control patients and patients with *S. aureus* respiratory tract infection (A), and in plasma from healthy volunteers and patients with *S. aureus* respiratory tract infection (B). The bars show mean ± SEM; samples from a prospective cohort. C) Plasma levels of CX_3_CL1 in *S. aureus* bacteremia (bloodstream infection) patients and healthy volunteers from a retrospective cohort. Values below limit of detection (LOD) was set to 0.1103 (*LOD*/√2) and are shown in grey. Statistical significances were calculated using Mann-Whitney Unpaired t-test (A-B) and Wilcoxon signed-rank test with median value compared to hypothetical value of 0.156 (i.e., in comparison with LOD) (C).

### *S. aureus*-associated CX_3_CL1 release is driven by α-toxin

As CX_3_CL1 levels were particularly high in patients with *S. aureus* respiratory tract infection, we performed follow up analyses using *in vitro* lung tissue models developed to study *S. aureus* pneumonia (6). The tissue models were stimulated with bacterial culture supernatants derived from the community-associated MRSA strain USA300 which has been associated with severe lung infections as well as two additional clinical *S. aureus* strains — strain NP796 isolated from a patient with a severe necrotizing pneumonia and strain LE2332 isolated from a patient with lung empyema (26). In the unstimulated lung model, CX_3_CL1 was evenly expressed throughout the epithelium, closely reflecting the CX_3_CL1 expression pattern in uninfected human lung tissue (Fig. 2 A). While unstimulated and LE2332 supernatant-stimulated lung models displayed similar patterns of CX_3_CL1 immunostaining, the lung models stimulated with the high toxin-producing strains USA300 or NP796 supernatants revealed a unique pattern of intensified CX_3_CL1 staining at the apical side of the epithelium compared to the deeper mucosal layers, which displayed relatively diffuse or no staining (Fig. 2 B). By analyzing the fluorescent ratio of CX_3_CL1 staining on the top (apical) and bottom (basal) half of the mucosal layer, we observed a significantly stronger apical expression of CX_3_CL1 in response to NP796 and USA300 supernatants as compared to that of the LE2332-treated and unstimulated models (Fig. 2 C). Importantly, these responses correlated with elevated levels of CX_3_CL1 in the lung model supernatants (Fig. 2 D). This suggests that the membrane-bound CX_3_CL1 is shed from the cell surface and released in response to *S. aureus* exotoxins.

**Figure 2.**
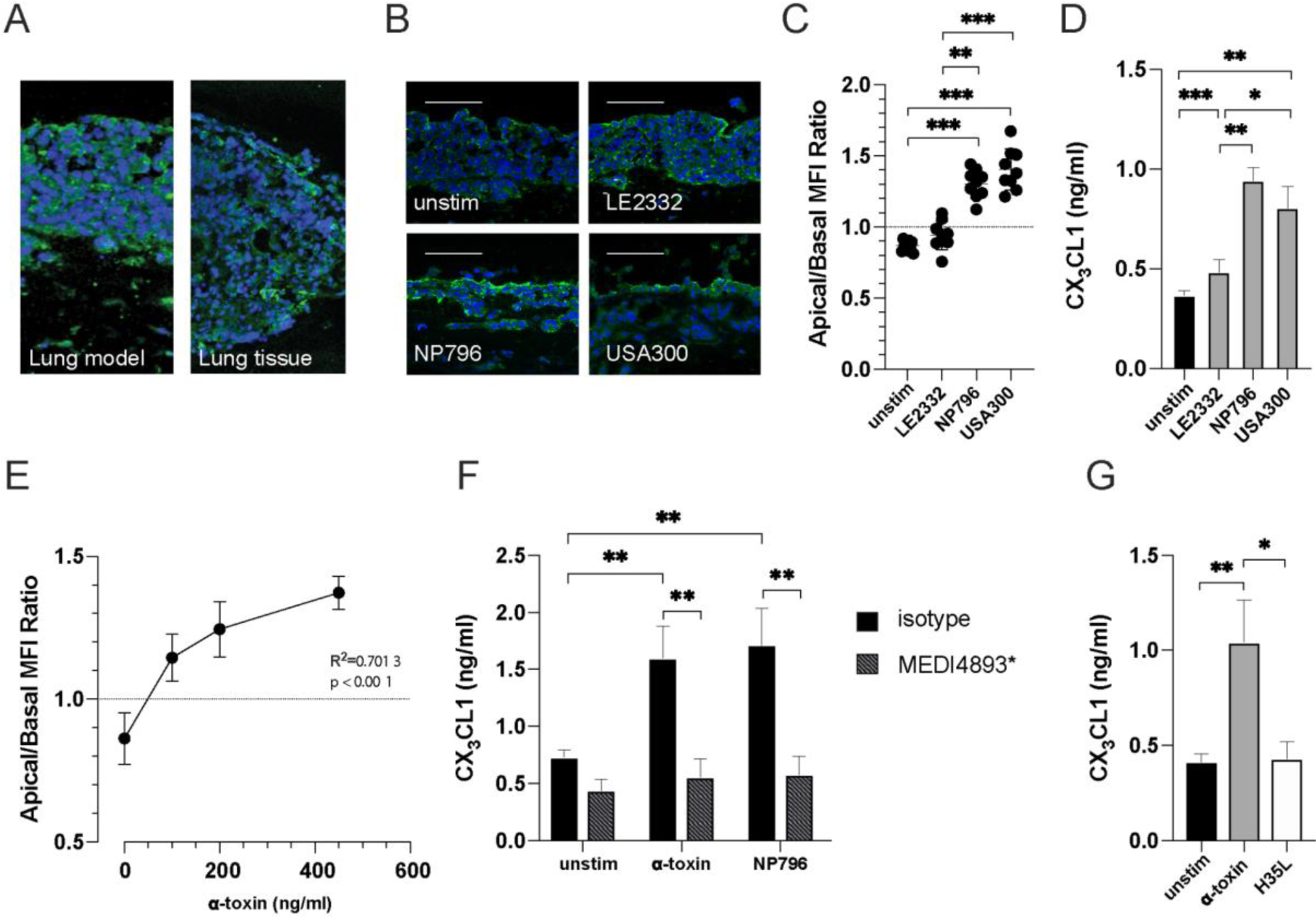
Alpha-toxin-mediated tissue redistribution and release of CX_3_CL1. A) Representative mean intensity projections of immunofluorescence microscopy images of tissue sections from lung tissue model and human lung tissue. The sections were stained with the nuclear stain 4′,6-diamidino-2-phenylindole, dihydrochloride (DAPI) (blue) in combination with antibodies detecting CX_3_CL1 (green). B) Immunofluorescence staining of CX_3_CL1 (green) and cell nuclei (blue, DAPI) in sectioned lung model stimulated with *S. aureus* LE2332, NP796, or USA300 bacterial culture supernatants (diluted 1:100). The scale bars in images equal 100 μm. C) Measurement of the spatial re-distribution (ratio of apical to basolateral mean fluorescence intensity, MFI) of CX_3_CL1 in lung tissue stimulated with bacterial culture supernatants (diluted 1:100). D) CX_3_CL1 levels in culture supernatants from unstimulated lung tissue model (black bar) or lung tissue models stimulated with LE2332, NP796, or USA300 bacterial culture supernatants (diluted 1:100) (grey). E) Measurement of the spatial re-distribution (ratio of apical to basolateral MFI) of CX_3_CL1 in lung models stimulated with various concentrations of α-toxin as indicated. F) CX_3_CL1 levels in culture supernatants of unstimulated lung models, or of lung models stimulated with α-toxin suspension (100 ng/ml) or NP796 bacterial culture supernatant (1:100) treated with anti-α-toxin antibodies (200 μg/ml, MEDI4893*) (striped bars). As control (black bar), an isotype matched antibody was used. G) CX_3_CL1 levels in culture supernatants of unstimulated lung model (black bars) or of lung models stimulated with wild type α-toxin (grey) or the mutated α-toxin H35L (white). ELISA determination of CX_3_CL1 levels was performed 24 h post stimulation. All experiments were performed at least three times. In D-G, the bars show the mean ± SD of three individual experiments. Statistically significant differences were determined by Kruskal-Wallis with uncorrected Dunn’s test in panel C-D and G, linear regression analysis in panel E, Ordinary two-way ANOVA with Sidak’s multiple comparison test in panel F.

An enhanced β-hemolytic phenotype on blood agar for USA300 and NP796 strains (Supplementary Fig 1 D) is consistent with the previously reported increased production of α-toxin in these strains as compared to LE2332 (26). An equally prominent β-hemolytic phenotype was found for *S. aureus* isolates from patients analyzed for CX_3_CL1 in lung airway fluids (Supplementary Fig 1 E). Thus, indicating that the above noted effect on CX_3_CL1 could be α-toxin mediated. Stimulation of lung models with increasing concentrations of α-toxin revealed a dose-dependent response in the apical/basal ratio of CX_3_CL1 expression in the lung models (R^2^ = 0.70, *p* < 0.001) (Fig. 2 E). Furthermore, addition of anti-α-toxin antibody (MEDI4893*) significantly mitigated the NP796 supernatant- and α-toxin-induced CX_3_CL1 release in the lung model supernatants (Fig. 2 F), suggesting that CX_3_CL1 released in *S. aureus* respiratory tract infection is driven by α-toxin. This is further supported by the reduced CX_3_CL1 levels in the BAL fluid of mice with staphylococcal respiratory tract infection that were administered the anti-α-toxin antibody MEDI4893* (Supplementary Fig. 2). Using H35L, a mutated α-toxin that is unable to form stable pores, we observed no change in the amount of soluble CX_3_CL1, suggesting the induction of CX_3_CL1 is dependent on α-toxińs pore-forming properties (Fig. 2 G).

### Alpha-toxin-induced CX_3_CL1 release is blocked by inhibiting ADAM10 activity

It has been suggested that the sheddase activity of ADAM10 can increase after calcium-influx as a consequence of α-toxin-mediated pore formation (17), thus leading to a loss of tissue integrity due to an increased rate of E-cadherin cleavage by ADAM10 (13). ADAM10 is also known to be responsible for shedding of membrane-bound CX_3_CL1 in steady state and during inflammation (Fig. 3 A) (22). Therefore, we investigated the role of ADAM10 in *S. aureus*-induced CX_3_CL1 release by using an inhibitor (GI254023X) of ADAM10 activity. Pretreatment of lung tissue models with GI254023X blocked the release of CX_3_CL1 induced by either α-toxin and NP796 supernatant (Fig. 3 B). In addition, in an *in vivo* murine model of *S. aureus* pneumonia we observed reduced CX_3_CL1 levels in the BAL fluid of mice treated with GI254023X prior to infection (Supplementary Fig. 3A). In the lung tissue models, GI254023X prevented also the altered expression pattern of CX_3_CL1 (Fig. 3 C-D). Furthermore, we demonstrated that blocking with anti-α-toxin antibody revealed a similar CX_3_CL1 expression pattern in the α-toxin treated lung model as that of the unstimulated model (Fig. 3 C-D). In addition, we analysed expression of E-cadherin, another factor known to be modulated by α-toxin mediated ADAM10 activity. The anti-α-toxin antibody blocking rescued loss of E-cadherin in the α-toxin treated lung model (Supplementary Fig. 3 B-C). While E-cadherin expression, as expected, disappeared from the tissue (Supplementary Fig. 3 BA-CB). CX3CL1 remained differently distributed (Fig. 3 C-D).

**Figure 3.**
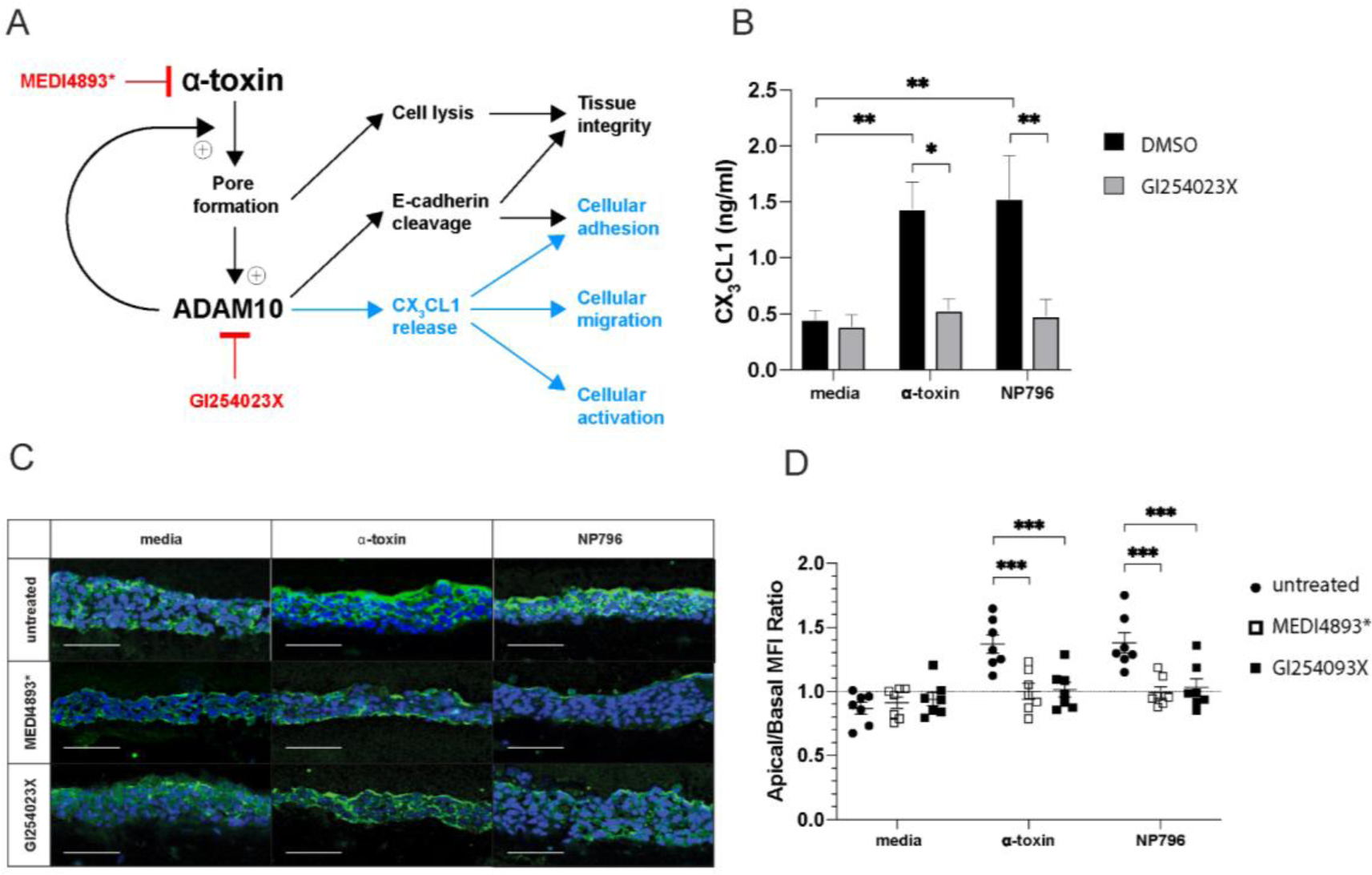
Alpha toxin-induced CX_3_CL1 release acts via ADAM10. A) Schematic drawing of the proposed pathway through which CX_3_CL1 is released from lung tissue exposed to α-toxin stimulation, and the proposed action of an anti-α-toxin antibody, MEDI4893*, or the ADAM10 inhibitor, GI254023X. B) CX_3_CL1 levels in culture supernatants of untreated lung models (black bars), or of lung models pre-treated with 10 µM of ADAM10 inhibitor, GI254023X (grey bars), 2 h post-treatment, the models were stimulated with α-toxin (100 ng/ml) or NP796 bacterial culture supernatant (1:100), or DMSO (GI254023X diluent). ELISA determination of CX_3_CL1 levels in supernatants of lung models was performed after 24 h of stimulation. C) Immunofluorescence staining for CX_3_CL1 (green) and cell nuclei (blue, DAPI) in sectioned lung model stimulated with *S. aureus* α-toxin (100 ng/ml) or NP796 bacterial culture supernatants (1:100), in the presence of either MEDI4893* (200 µg/mL) or GI254023X (10 µM) or left untreated. D) Measurement of the spatial re-distribution (ratio of apical to basolateral mean fluorescence intensity) of CX_3_CL1 in lung models stimulated with α-toxin suspensions (100 ng/ml) or NP796 bacterial culture supernatants (1:100), alone (untreated, circle) or treated with anti-α-toxin antibodies (200 μg/ml, MEDI4893*, open square), or the ADAM10 inhibitor, (10 µM, GI254023X, closed square). All experiments were performed at least three times. In B, the bars show the mean ± SD of three individual experiments. In C, the figure shows data from one representative experiment. In D, data are presented as the mean value ± SEM. The scale bars in images equal 100 μm. The statistical significances were calculated by ordinary two-way ANOVA, with Sidak’s multiple comparisons test in panel B and uncorrected Fisher’s LSD in panel D,

To assess whether ADAM10 metalloprotease enzyme activity and CX_3_CL1 release is a consequence of α-toxin mediated epithelial cell lysis (16HBE), we performed dose-response experiments where 16HBE cells were stimulated with increasing concentrations of α-toxin. Using LDH release assay, the cytotoxicity (15-10%) was weak at 100 ng/ml while it increased strongly at higher α-toxin concentrations (Supplementary Fig. 4A). Furthermore, ADAM10 metalloprotease enzyme activity was observed already at α-toxin concentrations from 50 ng /ml or higher (Supplementary Fig. 4B). Similarly, α-toxin-induced release of CX_3_CL1 was detected after stimulation with a concentration of 100 ng/ml or higher (Supplementary Fig, 4C). In all three assays, the α-toxin-mediated effects were inhibited with either the anti-α-toxin antibody MEDI4893* or the ADAM10 inhibitor GI254023X at all concentrations tested (Supplementary Fig. 4 A-C). These findings show that even lower α-toxin concentrations triggers ADAM10 activity and subsequent CX_3_CL1 release, and yet there is a clear link between increased α-toxin concentration and cytotoxicity, as well as ADAM10 activity and CX_3_CL1 release.

### Alpha-toxin-induced CX_3_CL1 release results in monocyte migration

Histological and immunohistochemical analysis of *S. aureus* infected lung tissue showed evidence of inflammation with substantial infiltration of immune cells, including CD14 positive cells (Fig. 4 A and Supplementary Fig. 5). Large amounts of bacteria were detected in lung tissue samples derived from *S. aureus*-infected patients with lung involvement as evident by a positive Brown-Brenn staining (Fig. 4A). Immunostaining for CD14 was prominent in areas with bacteria (Brown-Brenn positive) and coincided with cell-associated as well as more diffuse CX_3_CL1 staining patterns (Fig. 4A and Supplementary Fig. 5). CD14 and CX_3_CL1 were also relatively more prominent in the tissue of *S. aureus*-infected patients as compared to control tissue from individuals with other lung conditions (Fig 4A and B and Supplementary Fig. 5). As soluble CX_3_CL1 is a known chemotactic factor for monocytes (22), and to test whether the CX_3_CL1 shedding in the lung model coincides with enhanced migration of monocytes, we introduced blood monocytes into the lung models and monitored their motility with confocal live imaging following stimulation of models with α-toxin or NP796 supernatant. Both α-toxin and NP796 stimulation induced increased motility of monocyte-derived cells within the lung models (Fig. 5 A). To further test whether CX_3_CL1 released from stimulated lung models was responsible for the observed monocyte migration, we performed transwell migration assays using stimulated lung model supernatants on monocytes and neutrophils pre-treated with either a CX_3_CL1 receptor (CX_3_CR1) antagonist (AA-1-2008) or vehicle control. Both monocytes and neutrophils migrated in response to supernatants from α-toxin stimulated lung models; however, addition of the CX_3_CR1 antagonist (AA-1-2008) effectively reduced monocyte, but not neutrophil, migration to baseline levels (Fig. 5 B and Supplementary Fig. 6). Furthermore, CX_3_CL1 added to cell culture media induced CX_3_CR1-dependent monocyte migration (Fig. 5 B). However, we did not observe any CX_3_CR1-dependent migration of neutrophils given the same treatment (Supplementary Fig. 6). Together this suggests that *S. aureus* stimulation of lung tissue leads to the release of chemokines that attract both neutrophils and monocytes, with CX_3_CL1 as the major chemokine contributing to monocyte chemotaxis under these conditions.

**Figure 4.**
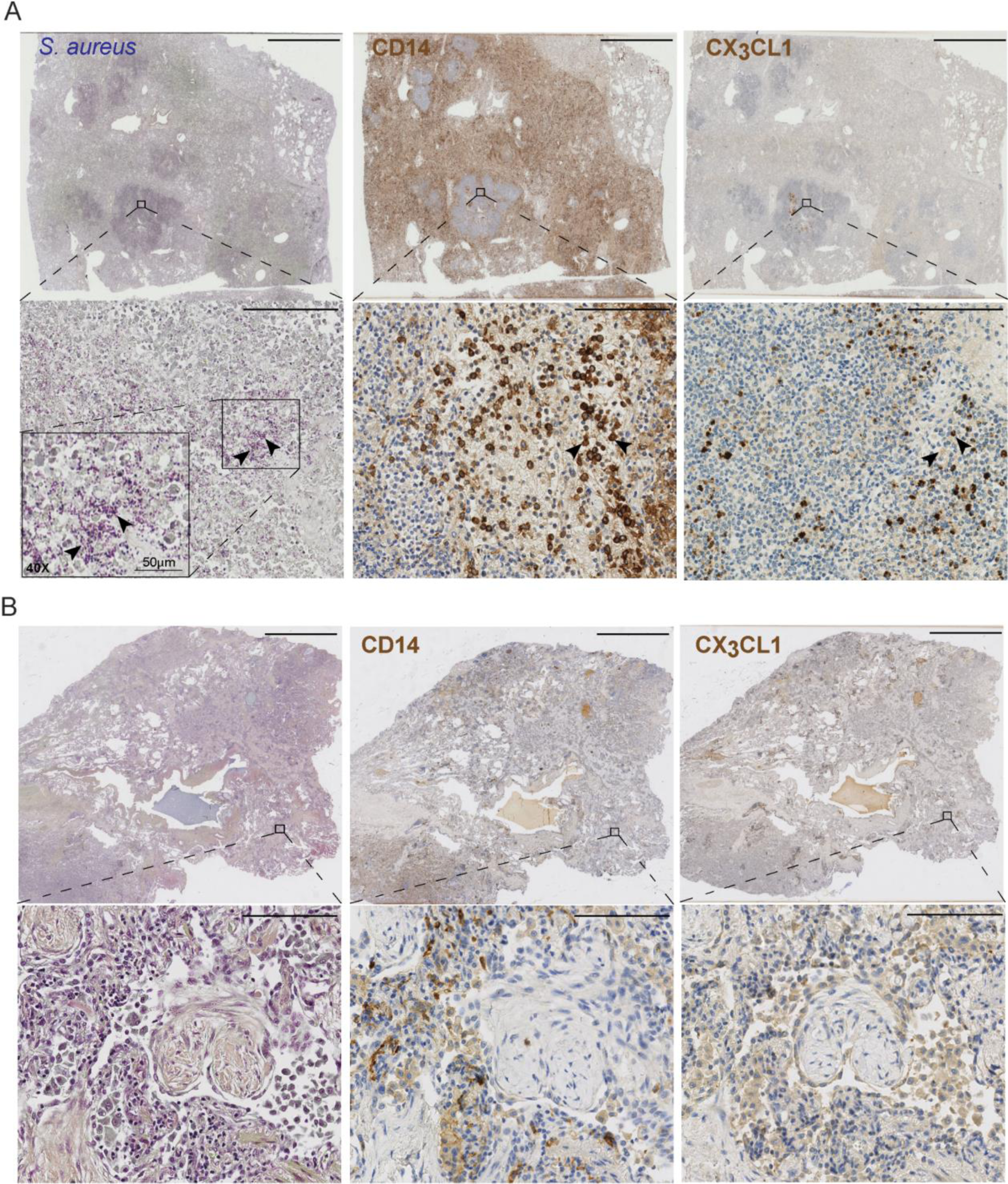
Histological and immunohistochemical qualitative analysis of *S. aureus* respiratory tract infection. Lung tissue biopsies were stained with Brown-Brenn reagents (Gram^+^ bacteria), anti-CD14 or anti-CX_3_CL1 antibodies. A) Immunohistochemical analysis for Gram^+^ *S. aureus* (Brown-Brenn, blue), CD14 (brown) and CX_3_CL1 (brown) of whole biopsy sections (top row) from an *S. aureus*-infected patient, with higher magnifications (20x, bottom row) of boxed areas. For the *S. aureus*-stained tissue a 40x magnification of one selected area in the 20x image is shown (insert). CD14 or CX_3_CL1 positive cells within this area of consecutive sections are indicated by arrows. The scale bars in images equal 5 mm (top row) or 100 μm (bottom row). B) Brown-Breen staining and immunohistochemical analysis for CD14 (brown) and CX_3_CL1 (brown) of whole biopsy sections (top row) from a non-infected fibrosis patient, with higher magnifications (20x, bottom row) of boxed areas. The scale bars in images equal 5 mm (top row) or 100 μm (bottom row).

**Figure 5.**
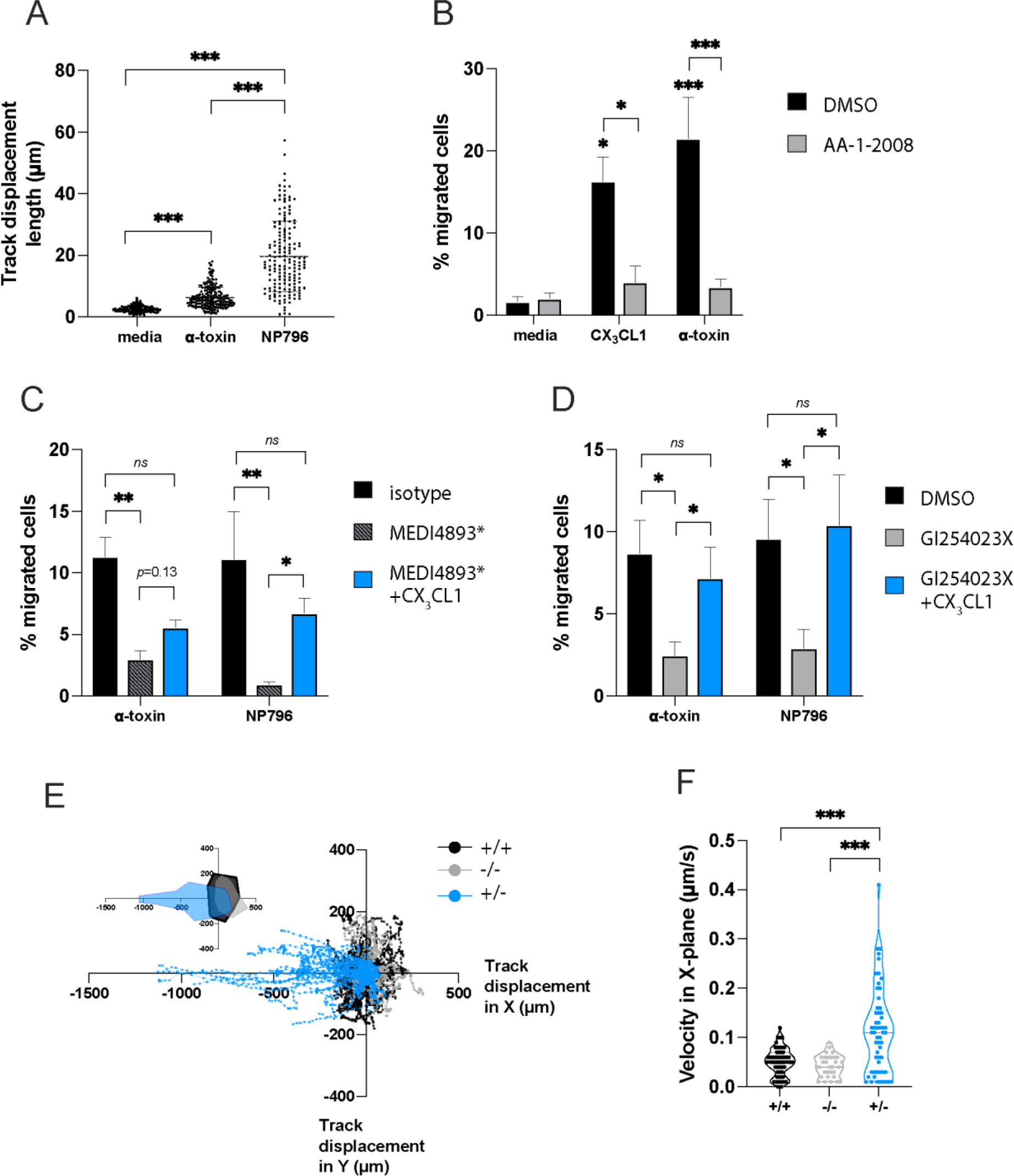
Monocyte migration in response to α-toxin-induced release of CX_3_CL1. A) Live imaging and total displacement determination of monocyte-derived macrophage-like cells in lung tissue models stimulated with α-toxin (100 ng/ml) or NP796 bacterial culture supernatant compared to unstimulated models (media) as indicated. B) Migration of human blood monocytes for 2 h in a transwell assay in response to supernatants from unstimulated lung model (media), lung models stimulated for 24-h with α-toxin (100 ng/ml), or medium supplemented with recombinant CX_3_CL1 (10 ng/ml), in the absence (DMSO, the diluent of AA-1-2008, black bars) or presence of the CX_3_CL1 receptor antagonist (AA-1-2008, 10 nM, grey bars). Pre-treatment with the CX_3_CR1 antagonist (AA-1-2008, 10nM) or DMSO (vehicle) was for 2 h prior to the migration assay. C) Migration of human blood monocytes for 2 h in a transwell assay in response to supernatants from lung models stimulated with α-toxin (100 ng/ml) or NP796 bacterial culture supernatant (1:100), treated with an isotype control-matched antibody (black bars, 200µg/ml) or anti-α-toxin antibodies (striped bars, 200 μg/ml, MEDI4893*). Supernatants from anti-α-toxin antibody treated conditions were supplemented with recombinant CX_3_CL1 (10 ng/ml, blue bars). D) Migration of human blood monocytes for 2 h in a transwell assay in response to supernatants from unstimulated lung models (black bars, DMSO GI254023X diluent), or of lung models pre-treated with the ADAM10 inhibitor (grey bars), GI254023X (10 µM) and then stimulated for 24 h with α-toxin (100 ng/ml) or NP796 bacterial culture supernatant (1:100), or medium from GI254023X-treated models supplemented with recombinant CX_3_CL1 (10 ng/ml, blue bars). E-F) Live imaging of monocyte directed migration towards a CX_3_CL1-gradient in a collagen matrix, total displacement (E) and velocity (F) for 2 h, is shown. In the migration chambers, CX_3_CL1 (10 ng/ml) was added on both sides (black) or excluded (grey) or added on one side (blue). Statistical significances were calculated by ordinary one-way ANOVA, with Tukey’s multiple comparisons test with a single pooled variance in panel A, C-D, ordinary two-way ANOVA, with Bonferroni’s multiple comparisons test in panel B, Kruskal-Wallis, with Dunn’s multiple comparisons test in panel F.

Supernatants from α-toxin- or NP796-stimulated lung models pre-treated with either anti-α-toxin neutralizing antibodies or an ADAM10 inhibitor failed to induce comparable levels of monocyte chemotaxis to the control supernatants (Fig. 5 C-D). The addition of exogenous CX_3_CL1 to supernatants from lung models pre-treated with ADAM10 inhibitor or anti-α-toxin antibodies restored the migratory capacity of the monocytes in the transwell migration assay (Fig. 5 C-D). To test whether CX_3_CL1 induce also directed monocyte migration in a collagen matrix, we performed 3D migration assays in a chamber that accommodates stable chemokine gradients for up to 48h. There was a clear preferential migration of monocytes towards CX_3_CL1 when added to one side of the µ-slide chamber (Fig. 5 E-F, Supplementary Video 1). These data suggest that soluble CX_3_CL1 released by α-toxin-stimulated lung models is both necessary and sufficient for the chemotaxis of human monocytes.

### CX_3_CL1 induces CD83 upregulation and impaired phagocytic killing capability of monocytes

To investigate *S. aureus*-induced responses beyond migration, we stimulated monocytes with lung model-derived supernatants containing CX_3_CL1 and analyzed expression of the co-stimulatory molecules CD83, CD86, and CD80 and the co-inhibitory molecule PD-L1, all of which are typically found on human myeloid cells. Monocytes were identified as described in Supplementary Fig. 7A and the analyses demonstrated that lung model supernatants stimulated with α-toxin or NP796 induced the upregulation of CD83 without a concomitant upregulation of CD86 or CD80 (Fig. 6 A, Band 6 D); all membrane proteins of the immunoglobulin receptor superfamily involved in the regulation of antigen presentation (27). The CD83 effect was reversed when α-toxin or ADAM10 were blocked prior to stimulation of the lung model (Fig. 6A). Upregulation of CD83 was also seen when CX_3_CL1 was added to unstimulated lung model supernatants, suggesting CX_3_CL1-induced alteration of co-stimulatory molecules on monocytes (Fig. 6 A). Alpha-toxin stimulated lung model supernatants also induced a down regulation of CD86 as well an upregulation of PD-L1 (Fig. 6 B and C), a protein belonging to the family of immune checkpoint molecules and a co-inhibitory receptor regulating T cell activation. The alterations in CD86 and PD-L1 surface expression were reversed when α-toxin or ADAM10 was blocked during stimulation with supernatants from α-toxin-stimulated lung models (Fig 6 B and C). Unstimulated lung model supernatants spiked with CX_3_CL1 or NP796-stimulated lung model supernatant had no detectable effect on PD-L1 (Fig. 6 C). Furthermore, the changes in CD80 expression observed with NP796-stimulated lung model supernatants could not be reproduced with CX_3_CL1-spiked lung model supernatants (Fig. 6 D).

**Figure 6.**
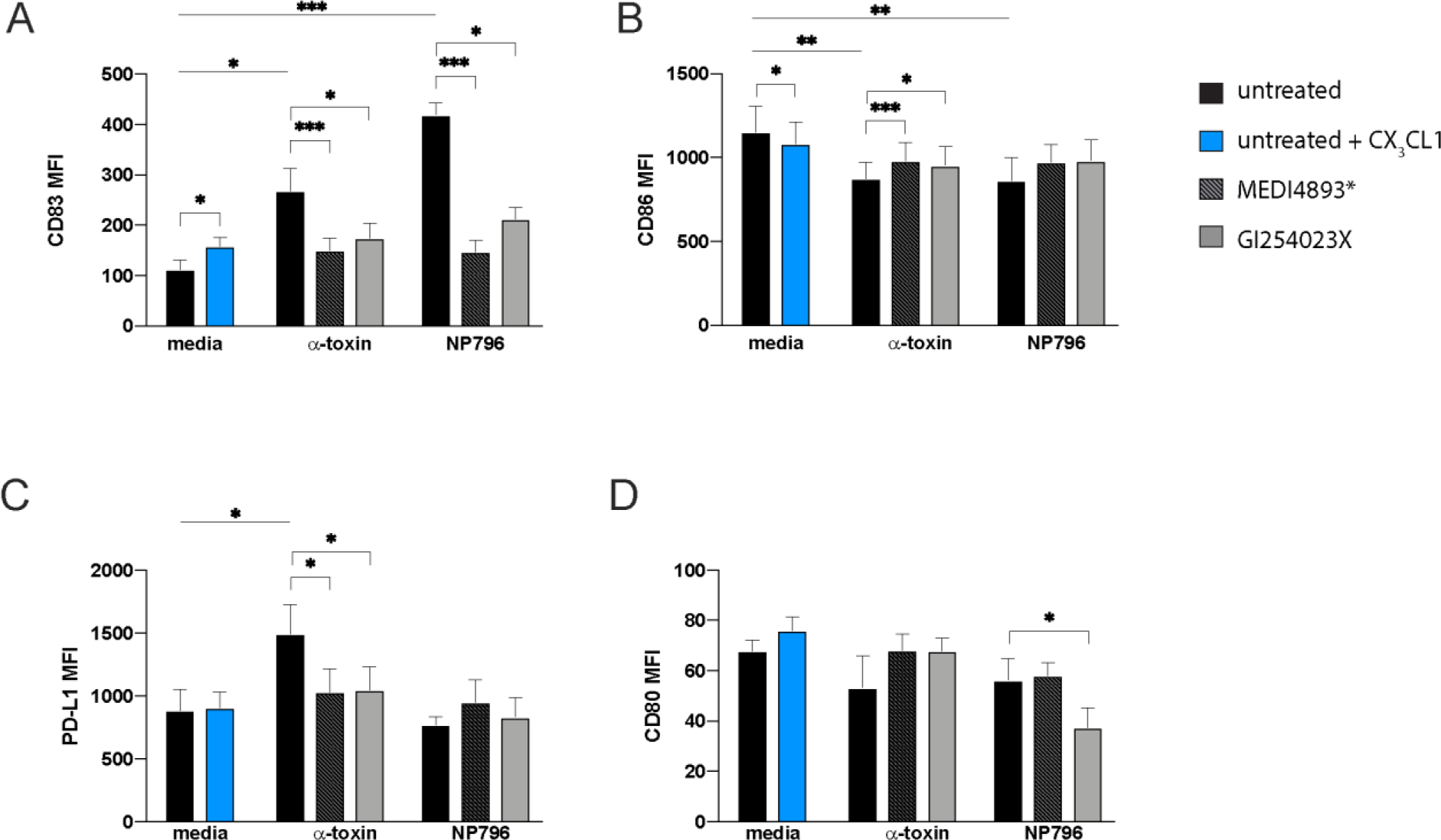
CX_3_CL1 modulates expression of co-stimulatory/inhibitory molecules. A-D) Monocytes were stimulated or left unstimulated for 24 h and subsequently processed for flow cytometry analysis. MFI for CD83 (A), CD86 (B), PD-L1 (C) and CD80 (D) on HLA-DR^+^ CD14^+^ monocytes cultured with supernatants from unstimulated lung models (black bars, media), or unstimulated lung model supernatants spiked with 10ng/ml CX_3_CL1 (blue bars), or lung models stimulated with α-toxin (100 ng/ml) or NP796 bacterial culture supernatant (1:100) (black bars) as indicated. For CX_3_CL1 stimulation of monocytes, CX_3_CL1 was added at 10ng/ml to unstimulated lung model supernatants (blue). Lung model supernatants used to stimulate monocytes were also generated by treating the α-toxin suspension or NP796 supernatant with anti-α-toxin antibodies (striped bars, 200 μg/ml, MEDI4893*), or by treating the lung model with the ADAM10 inhibitor (grey bars, GI254023X, 10 µM) prior to stimulation with either α-toxin or NP796 supernatant. The bars show the mean ± SEM of 8 independent donors, and statistical significances were calculated by Wilcoxon signed-rank test and Friedman test, uncorrected Dunn’s test.

To study the effects of CX_3_CL1 on monocytic anti-microbial capabilities, we infected CX_3_CL1-pretreated monocytes with *S. aureus* and quantified phagocyic uptake as well as killing of intracellular bacteria. We used the two *S. aureus* strains, Cowan I and LE2332, instead of NP796 since this strain caused rapid lysis of the monocytes. Intracellular bacteria were identified using imaging flow cytometry (ImageStreamX) and a GFP-expressing Cowan I strain (Fig. 7 A). We designated bacteria residing inside the monocytes after one hour as phagocytosed. The intracellular killing of GFP-producing Cowan I was estimated by calculating the percentage of bacteria that were detectable 4 h post infection. Monocytes pre-treated with CX_3_CL1 exhibited a decreased ability to kill intracellular bacteria, while their ability to phagocytose remained unchanged (Fig. 7 B). To verify that GFP-signal detected by imaging flow cytometry represented live bacteria, we lysed the monocytes and plated the bacteria. The data confirmed similar uptake by monocytes and reduced intracellular bacterial killing by monocytes pre-treated with CX_3_CL1 (Fig. 7 C). Pre-treatment with lung model supernatants from NP796-stimulated models yielded similar results to CX_3_CL1-pretreated monocytes with respect to CFU counts, and the effect was abrogated by blocking α-toxin or ADAM10 prior to stimulation of the models (Fig. 7 D). The decreased ability to kill phagocytosed bacteria in the presence of CX_3_CL1 was consistent with a decrease in both ROS and NO production (Fig. 7 E-F and Supplementary Fig 8). Taken together, we observed phenotypic and functional changes that suggest impaired monocyte function following exposure of CX_3_CL1 as a consequence of ADAM10-mediated activity induced by α-toxin.

**Figure 7.**
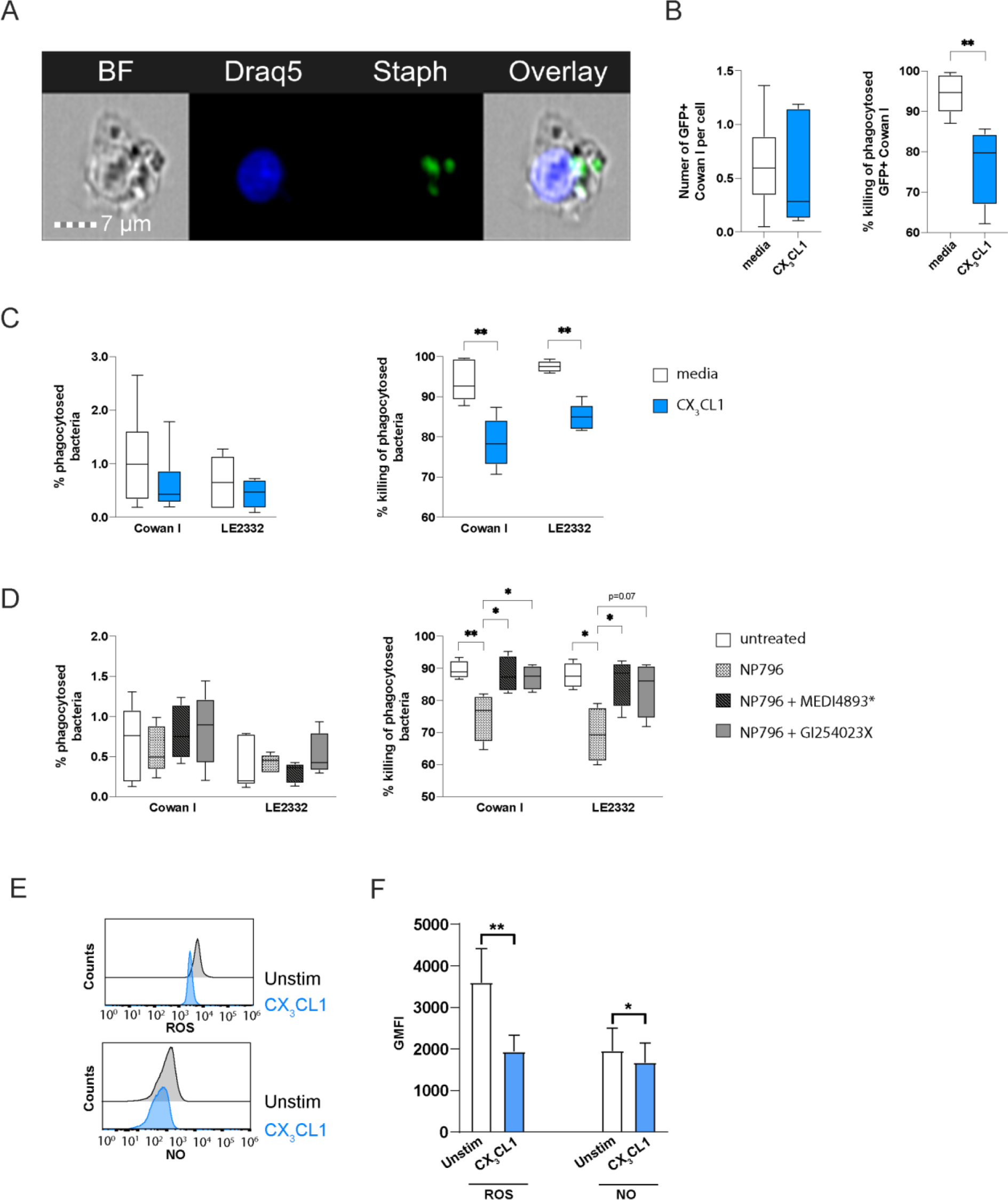
CX_3_CL1 modulates monocyte function. A-C) Monocytes stimulated with CX_3_CL1 (10 ng/ml) or left unstimulated, were infected with *S. aureus* Cowan I-GFP, fixed and stained with Draq5, and analysed by imaging flow cytometry or CFU assay. A) Representative images showing monocytes containing bacteria. B) Image flow quantification of the proportion of Cowan I-GFP^+^ monocytes after 1 h (left) and after 4 h (right). C) Percentage phagocytosed Cowan I or LE2332 (1h, left) and percentage killing of phagocytosed Cowan I or LE2332 (4h, right) after stimulation with media (white), CX_3_CL1 (10 ng/ml, blue). The bars show the mean ± SEM of four independent donors. D) Percentage phagocytosed Cowan I or LE2332 (1h, left) and percentage killing of phagocytosed Cowan I or LE2332 (4h, right) after stimulation with lung model supernatants as indicated. The lung model supernatants were obtained by stimulating lung models for 24 h with NP796 bacterial culture supernatant alone (checkered, 1:100), NP796 bacterial culture supernatant treated with MEDI4893* (grey striped, 200µg/ml), or NP796 bacterial culture supernatant in the presence of GI254023X (grey, 10 µM). E-F) Monocytes stimulated with CX_3_CL1 (10 ng/ml) for 18 h or left unstimulated, were infected with *S. aureus* strain LE2332 (MOI 1) and analyzed for ROS and NO production. Representative histograms (E) of ROS and NO production, and representative bar graphs (F) of ROS and NO production by LE2332-infected monocytes. The bar graphs show the mean ± SEM of five independent donors. In B-D, data presented as whisker plots with box indicating interquartile range and error bars indicating highest and lower value of four independent donors, and statistical significances were calculated by Mann-Whitney test in B and C and Kruskal-Wallis with uncorrected Dunn’s test in D. In F statistical significances were calculated by Wilcoxon matched-pair signed-rank test.

## DISCUSSION

In this study, we investigated the mechanism and consequences of *S. aureus*-associated release of CX_3_CL1, a unique membrane-bound chemokine that requires metalloprotease processing before secretion. We found that CX_3_CL1 levels increase both locally and systemically in patients with *S. aureus* respiratory tract infection. Using a 3D human *in vitro* staphylococcal pneumonia model, we show that α-toxin-induced activation of ADAM10 leads to increased CX_3_CL1 release, which enhanced monocyte motility and impaired intracellular killing capacity by monocytes. The reduced bactericidal phenotype was accompanied by an increased CD83 and a decreased CD86 surface expression. The results imply that α-toxin alters the inflammatory milieu by which CX_3_CL1 affects immune cell functions beyond cellular adhesion and migration that contributes to *S. aureus* intracellular survival and pathogenesis.

*S. aureus* toxins interact with host tissue constituent cells as well as immune cells. To fully appreciate the role of *S. aureus* toxins in human disease, studies of *S. aureus* infections have to take into consideration the host-specificities as well as the cellular and molecular interactions occurring within the relevant tissue. In this study, we found elevated levels of CX_3_CL1 in patients with *S. aureus* infections—especially in infections involving the respiratory tract. Exploiting the human lung model enabled us to study the mechanisms by which *S. aureus* triggers CX_3_CL1 release in human-derived lung epithelial cells. The blocking effect of the anti-α-toxin specific antibody demonstrated that it was specifically α-toxin among the various *S. aureus* toxins that led to the redistribution and release of CX_3_CL1 in the lung model. Furthermore, using a mutated α-toxin variant our results support the dependency of α-toxin’s pore-forming activity in the activation and release of CX_3_CL1 from the membrane. In addition to α-toxin, other *S. aureus* pore-forming toxins, e.g., Panton-Valentine Lekocidin (PVL) and γ-hemolysins, may also affect signaling pathways in the respiratory tract (28–30). However, the availability of receptors for the *S. aureus* pore forming toxins can explain cellular and tissue tropism; ADAM10 is highly expressed on epithelial cells while the receptors for PVL and γ-hemolysins are predominantly found on human myeloid cells. Whether the different pore-forming toxins exhibit coordinated activities to thwart host defenses and establish a successful infection needs to be investigated further. To encompass the potential effects of various cells on tissue responses during toxin exposure, models with extended cell complexity need to be developed. This is particularly relevant considering that these toxins, initially recognized for their ability to disrupt eukaryotic membrane barriers and cause target host cell lysis, are now understood to also exert subtle changes in cell activity and host physiology, even at sub-lytic concentrations (9, 31, 32). This notion is in line with our results demonstrating that although there is a clear association between degree of α-toxin mediated cytotoxicity and CX3CL1 release, the release was noted even at sub-lytic toxin concentrations.

The release of CX_3_CL1 can be mediated by several different metalloproteinases including ADAM10 and ADAM17 (22, 33–35), and here we show for the first time that this ADAM-10-CX_3_CL1 axis is triggered by *S. aureus* virulence factors. As ADAM10 is known to cleave membrane-bound CX_3_CL1 and also serve as the main receptor for α-toxin, we examined the effect of the ADAM10-inhibitor GI254023X on α-toxin induced CX_3_CL1 release. We observed that both GI254023X and the anti-α-toxin antibody blocked the release of CX_3_CL1 in our toxin-stimulated *in vitro* model. Since ADAM10 and ADAM17 share substrates, including CX_3_CL1 (36), we cannot rule out the involvement of ADAM17 in the release of CX_3_CL1. However, GI254023X has a thousand-fold higher affinity for ADAM10 and is generally regarded as an ADAM10-specific inhibitor. Therefore, the effect mediated by GI254023X should, to a high degree, be attributed to loss of ADAM10 activity.

Although it cannot be excluded that freezing as well as the timing of plasma collection may impact on the levels of CX_3_CL1 detected, the data indicate that *S. aureus* infection with lung involvement leads to particularly high CX_3_CL1 levels both locally and systemically. In line with the reasoning of a lung-infection contribution, we observed increased CX_3_CL1 levels in patients with confirmed *S. aureus* respiratory infection while a large number of plasma samples from our retrospective cohort of critically ill patients with bloodstream *S. aureus* infection but without respiratory tract involvement had undetectable levels of CX_3_CL1. We did not find an association with co-morbidities or with severity of disease. This observation is in contrast to a study by Pachot *et al*. in which a population of critically ill patients with septic shock was examined and where an increase in CX_3_CL1 and positive correlation with disease severity was observed (37). In this context, it is interesting that as many as 50 % of the patients included in the study by Pachot *et al*., had pulmonary infection listed as the primary site of infection. Because the study by Pachot and colleagues did not take aetiology into account, direct comparisons must be made with caution. However, the high percentage of lung focus in that study is in line with our data suggesting that the measured increase in CX_3_CL1 might be particularly pronounced due to respiratory tract infection caused by *S. aureus*.

Alpha-toxin-stimulated lung models induced migration in both monocytes and neutrophils. In this context, CX_3_CL1 was one of the most significant chemokines for monocyte migration, as evident by the finding that CX_3_CR1 receptor inhibition decreased monocyte migration to base line levels. Inhibiting CX_3_CR1, on the other hand, had no effect on neutrophil migration, suggesting that other chemokines released from lung tissue are important for neutrophil migration in *S. aureus* infection. The independence of neutrophil migration from CX_3_CL1 can probably be explained by CXCL8, a prototypical neutrophil chemoattractant, released from *S. aureus* and α-toxin-stimulated lung models (6). That CX_3_CL1 plays a minor role in human neutrophil migration was further strengthen by the observation that addition of CX_3_CL1 did not act as a chemoattractant for neutrophils. This is also in line with the fact that neutrophils express relatively low of levels of CX_3_CR1 (38), while human monocytes are highly positive for CX_3_CR1 (39).

In addition, we observed that CX_3_CL1 induced phenotypic and functional changes in monocytes including an upregulation of CD83 expression. Although often used as a marker for activated dendritic cells (DCs), soluble CD83 has been shown to interfere with (DC)-induced T cell stimulation (40). It has also been suggested that monocytes, in contrast to DCs, produce and release CD83 which may have immune suppressive functions by interacting with membrane-bound CD83 on activated DCs (41, 42). In further support of a suppressive phenotype, we also observed a reduction in the activation marker, CD86, on the surface of CX3CL1-treated monocytes. Notably, the CX_3_CL1-treated monocytes revealed a reduced capacity to kill phagocytosed *S. aureus*. This functional impairment was linked to a reduction of both NO and ROS. These effector molecules have a direct antimicrobial effect, and the products generated during the respiratory burst may also act as second messengers in redox signaling and cellular activation processes (43, 44).

It is known that CX_3_CL1 is constitutively expressed in epithelial and endothelial tissues and has functions related to cell anchoring and adhesion (39). Furthermore, ADAM10 constitutively sheds CX_3_CL1 at low levels (22) which strongly suggests that ADAM10 and CX_3_CL1 have functions related to maintaining tissue homeostasis. Our observations of released CX_3_CL1 and its effects on monocytes after α-toxin stimulation suggest that *S. aureus* is exploiting CX_3_CL1’s homeostatic functions to promote bacterial survival. There is accumulating evidence that the CX_3_CL1-CX_3_CR1 axis is involved in various diseases, including atherosclerosis (25) and rheumatoid arthritis (45), as well as pulmonary disorders (46, 47). In the respiratory tract, CX_3_CL1 has also been proposed to support the migration of monocytes in interstitial lung disease (48) and to promote pulmonary fibrosis by attracting macrophages with a pro-fibrotic phenotype (49). The CX_3_CL1-CX_3_CR1 axis is also suggested to be involved in the pathogenesis of critically ill and septic shock patients as evident by a link between reduced CX_3_CR1 mRNA expression in circulating leukocytes and increased mortality (24, 37). Our study is the first to show that *S. aureus* induces shedding of CX_3_CL1 by lung epithelial cells via α-toxin-mediated activation of ADAM10, with the potential to modulate immune responses locally and impair monocyte’s antimicrobial function thereby contributing towards *S. aureus* pathogenesis and survival during respiratory infections.

## MATERIALS AND METHODS

### Patients and samples

Lung tissue, bronchioalveolar lavage (BAL), tracheal aspirates and blood of patients were collected from patients at the University Hospital Zurich, Switzerland, after obtaining informed consent. All blood and tracheal aspirate samples were processed and immediately analyzed for CX_3_CL1. Collection of patient samples were done in accordance with the Helsinki declaration and performed with permission from the Regional Ethics Board in Zurich, Switzerland.

Patients with *S. aureus* bacteremia were recruited at the Department of Infectious Diseases, Örebro University Hospital, Sweden, after signed informed consents were obtained. The demographic data together with the comorbidities are provided in Supplementary Table 1. Sepsis was defined according to the Sepsis-3 definition. Blood samples were collected one or two days after hospital admission, i.e., when blood culture signaled positive, and further processed for collection of plasma for freezing and storage at −80°C. The study was done in accordance with the Helsinki declaration and performed with permission from the Regional Ethics Board of Uppsala, Sweden.

Blood samples from healthy volunteers (controls) were collected in University Hospital Zurich or in the Department of Clinical Immunology and Transfusion Medicine at the Karolinska University Hospital and were further processed for isolation of monocytes and collection of plasma. Monocytes were analyzed fresh and plasma were analyzed fresh or after freezing and storage in −80°C. All healthy volunteers were included after providing signed informed consent. Blood collections were performed in accordance with the Helsinki declaration and with permission from the Regional Ethics Board, in Stockholm, Sweden or Cantonal Ethics Board, in Zurich, Switzerland.

### Bacterial Strains

Clinical *S. aureus* isolates used in this study were collected from the pleural fluid of a necrotizing pneumonia case (NP796) and one lung empyema case (LE2332) at Global Hospitals, Hyderabad, India (6). USA300 (11358) (50) and Cowan I (ATCC 12598) (51) was used as reference strains. In addition, four clinical *S. aureus* strains (1408, 1439, 1441, and 1525) from patients with *S. aureus* respiratory tract infection were used in the hemolysis assay. *S. aureus* Cowan I (ATCC 12598) was transformed with the reporter plasmid pCN56 encoding the fluorescence-activated cell sorting-optimized mutant version gfpmut2 of the green fluorescent gene of *A. victoria* encoding the GFP reporter and a constitutive *blaZ* promotor (52). The blaZ promotor was integrated into the multiple cloning site of pCN56 by using the restriction enzymes SphI and BamH1. Before transformation of *S. aureus* Cowan I, the plasmid was multiplied in *E. coli* IM30B (53) to bypass restriction systems.

### Lung tissue model set up

The lung tissue model was set up as previously described (54). Briefly, lung tissue models were established using 8×10^4^ MRC-5 cells (human lung fibroblast cell line) at passage < 30 cultured in a collagen matrix for seven days, followed by addition of 1.2 x 10^5^ 16HBE14o-cells (human bronchial epithelial cell line) per model. Epithelial cells were confluent after 3 days and were then air-exposed for 7 days. Complete DMEM (DMEM supplemented with 10% heat-inactivated fetal calf serum, 2mM L-glutamine, 10 mM HEPES) was changed every 2-3 days. Lung tissue model were exposed to stimuli in a 50µl droplet on the apical side. For models containing monocyte-derived cells, monocytes were isolated as described, stained with PKH26 Red Fluorescent Cell Linker Kit for General Cell Membrane (Sigma-Aldrich, Saint Louis, Missouri, USA) according to manufacturer’s instructions and added at a concentration of 5×10^5^ monocytes per model the day before adding epithelial cells.

### Isolation of monocytes and neutrophils

Human peripheral blood mononuclear cells (PBMCs) were isolated from buffy-coated or whole blood from healthy volunteers, using Lymphoprep (Axis-Shield, Oslo, Norway) gradient centrifugation according to the manufacturers’ instructions. Monocytes were isolated from PBMCs using EasySep Human monocyte enrichment kit without CD16 depletion (StemCell Technologies, Vancouver, British Columbia, Canada) according to the manufacturers’ instructions. Human neutrophils were isolated from fresh whole blood obtained from healthy volunteers using Polymorphprep (Axis-Shield, Oslo, Norway) by performing density gradient separation according to the manufacturers’ instructions.

### Hemolysis assay

For the hemolysis assays, human red blood cells (RBCs) were isolated from heparinized venous blood obtained from healthy adult volunteers. RBCs were washed twice in sterile saline (0.9% NaCl) and centrifuged at 500 x g for 10 min. RBCs were diluted to 5% in PBS. 100 μL of RBCs was incubated with 100 μL of bacterial culture supernatant in a 96 well plate for 30 min at 37 °C. A dilution series of bacterial culture supernatant was prepared using PBS with 1:1 being the highest concentration. Plates were centrifuged for 5 min at 500 x g and supernatants were transferred to a sterile 96-well plate and RBC lysis evaluated by determining the absorbance at 415nm. PBS alone and H_2_O were used as negative and positive controls, respectively.

### Bacterial supernatant preparation

Bacterial culture supernatants for the stimulation experiments were prepared as previously described (26). In short, the strains were cultured overnight at 37°C in 25 ml casein hydrolysate and yeast extract (CCY) medium. Cell-free supernatants were prepared through centrifugation at 3350 g followed by filter sterilization.

### Lung tissue model stimulation

Once established, lung tissue models were stimulated with 1:100 dilutions of bacterial supernatants in PBS or cell culture media. Antibody blocking of α-toxin with MEDI4893* was performed by pre-incubating the bacterial culture supernatants for 1 h before stimulating the tissue models. Similarly, tissue models were pretreated with ADAM10 inhibitor (GI254023X) for 2 h at 37°C before stimulations. All stimulations, and treatment of tissue model prior to stimulation, were done by adding a 50µl drop on the apical side of the model. Prior to cryosectioning, models were submerged in 2M sucrose solution for 90-120 minutes, followed by cutting out and embedding in O.C.T. (Sakura Finetek Europe, Zoeterwoude, Netherlands) and subsequent frozen at −20°C for 24 h and then transferred to −80°C until used. The cell lines MRC5 and 16HBE14o-were routinely tested for mycoplasma contamination.

### Immunostaining and confocal microscopy

Cryosectioning of lung tissue models were performed as previously described (6). Briefly, 10 μm sections were obtained using a MICROM cryostat HM 560 MV (Carl Zeiss, Oberkochen, Germany) and fixed in 2% freshly prepared paraformaldehyde in PBS for 15 min at room temperature. Immunofluorescence staining of lung tissue model sections for confocal microscopy (54) were performed as previously described. Mouse anti-E-cadherin (2 μg/ml, clone HECD-1; Invitrogen, Carlsbad, California, USA) and mouse anti-CX_3_CL1 (2 μg/ml, clone MM0207-8J23; Abcam, Cambridge, United Kingdom) were used to stain the sections of lung tissue models. Specific staining was detected by Alexa Fluor 488-conjugated donkey anti-mouse IgG (3.3 μg/ml) (Molecular Probes, Eugene, Oregon, USA). Staining was visualized using a Nikon A1 confocal microscope (Nikon Instruments, Tokyo, Japan). The mean fluorescence intensity (MFI for E-cadherin) and Apical/Basal MFI ratio (for CX_3_CL_1_) in 4-5 fields per tissue section was determined using Image J analysis software (Fiji).

### Histology and immunohistochemistry

Patient tissue were fixed, sectioned and stained as described as described previously (55). Patient tissues were first fixed in 4% buffered formalin and then paraffin-embedded (Leica, Muttenz, Switzerland). Two μm sections were stained with Brown-Brenn to visualize bacteria in the tissue. Immunohistochemistry staining was used to detect human CD14 and human CX_3_CL1 (Abcam; MM0207-8J23) on paraffin-embedded biopsies. Whole-slide scanning and photomicrography were performed with a NanoZoomer 2.0-HT digital slide Scanner (Hamamatsu, Houston, TX).

### ELISA

The levels of CX_3_CL1 were determined according to manufacturer’s instructions using the Human CX_3_CL1/Fractalkine DuoSet ELISA and DuoSet ELISA Ancillary Reagent Kit (R&D Systems, Minneapolis, Minnesota, USA). Supernatants were diluted 1:1 with Reagent Diluent for all measurements and performed in duplicates. For CX_3_CL1 the limit of detection was 0.156 ng/ml. Values below LOD was set to 0.1103 (*LOD*/√2). The levels of CD83 (pg/ml) secreted in the culture medium were determined with the Human CD83 DuoSet ELISA kit (R&D Systems, Minneapolis, Minnesota, USA) according to manufacturer’s instructions. Supernatants were analyzed undiluted and all measurements were performed in duplicates.

### *In vivo* blocking of α-toxin and ADAM10

Blocking of α-toxin in an experimental *in vivo* model *S. aureus* respiratory tract infection was performed as described in supplementary online information (Supplementary File – Text). *S. aureus* frozen stock cultures were thawed and diluted to the appropriate inoculum in sterile PBS, pH 7.2 (Invitrogen) (56). Specific-pathogen-free 7- to 8-week-old female C57BL/6J mice (The Jackson Laboratory, Bar Harbor, ME) were briefly anesthetized and maintained in 3% isoflurane (Butler Schein™ Animal Health) with oxygen at 3 L/min and infected intranasally. Anti α-toxin MEDI4893* or c-IgG was administered in 0.5 mL intraperitoneally (IP) 24h prior to infection. Animals were euthanized with CO_2_ 24h post infection, and BAL fluid was collected for CX_3_CL1 measurement. All animal studies were approved by the AstraZeneca Institutional Animal Care and Use Committee and were conducted in an Association for Accreditation and Assessment Laboratory Animal Care (AAALAC)-accredited facility in compliance with U.S. regulations governing the housing and use of animals. 7- to 9-week-old female C57BL/6J mice (Janvier Labs, France) were pre-treated with GI254023X diluted in 0.1 M carbonate buffer to a 0.014 M stock, and ∼200 mg kg-1 or PBS with carbonate buffer as control in a final volume of 100 ul via intraperitoneal injection. Pre-treatment was applied daily for 3 to 5 days prior to the infection with *S. aureus*. On the day of the infection, mice were anesthetized with ketamine/xylazine injected intraperitoneally (ketamine 65-90 mg/kg and xylazine 10-13 mg/kg). The anesthetized mice were infected intranasally with the *S. aureus* strain USA300 (5 x 10^7^ CFU). At the end point the mice were euthanized with CO_2_ and BAL fluid was collected by washing the lungs with PBS for CX_3_CL1 measurement. The protocols (ZH050/18) were approved by the institutional animal care and use committee of the University of Zurich and all experiments were conducted in approval of the Cantonal Veterinary Office Zurich.

### ADAM10 metalloprotease enzymatic activity assay

ADAM10 enzymatic activity was measured on the basis of cleavage of fluorescent substrate using the Mca-P-L-A-Q-A-V-Dpa-R-S-S-S-R-NH2 Fluorogenic Peptide Substrate III (R & D systems). 16HBE cells were seeded at 50,000 cells per well in a 96 well plate. The cells were washed and stimulated with 10, 50, 100, 250, 500 1000 ng/ml of α-toxin for 2h in the absence or presence of anti-α-toxin antibodies (200 μg/ml, MEDI4893*) or the ADAM10 inhibitor, GI254023X (10 µM). 16HBE cells were pre-treated with GI254023X for 4h or α-toxin was preincubated with MEDI4893* for 4h before stimulations. The ADAM10 enzymatic activity was determined by incubating the cells in 25 mM Tris buffer at a pH 8.0 with 10 μM with fluorogenic peptide substrate for 30 min at 37°C in a 100 μl final reaction volume. Fluorescence intensity was read on a SpectraMax iD3, Molecular Devices plate reader.

### Lactate dehydrogenase (LDH) activity assay

Cytotoxic responses were assessed by measuring LDH release into the tissue culture medium by cells that had been stimulated with bacterial culture supernatants or pure toxins. LDH was measured using the Promega CytoTox 96 Nonradioactive Cytotoxicity assay kit according to the manufacturer’s protocol. The absorbance was read at 490 nm using a SpectraMax iD3, Molecular Devices plate reader. Percent cytotoxicity was determined in relation to the lysis control.

### Live Imaging

For live imaging, models were prepared as previously described (57). Briefly, models were kept for 4 days post air-exposure and fixed upside-down on a 6-well No. 0 Coverslip 10 mm Glass Diameter uncoated plate (MatTek, Ashland, Massachusetts, USA). Models were kept in a humidified CO_2_ chamber during the whole imaging process, and images were taken every 20 minutes for approximately 16 h. For each model, a z-stack of approximately 120 µm around the epithelial layer was acquired with a 547 nm laser. Image processing and analysis of reconstructed three-dimensional Z-stack were done using Imaris (Bitplane, Andor Technology, Belfast, United Kingdom). Cells were tracked using identical settings across all stimulations with the ImarisTrack module.

### Transwell migration assay

The chemotactic effect of CX3CL1 (10 ng/ml) and lung conditioned media (supernatants collected from stimulated lung tissue models) were assessed using a Transwell migration assay. The lung conditioned media (diluted 1:20 in RPMI 1640 media) were added to the outer chamber of 24-well transwell plates (3 µm Costar, Corning Inc, New York, USA). Neutrophils and monocytes were seeded at 3 x 10^5^ to 5 x 10^5^ cells/well in the upper chamber of a 24-well Transwell plate and incubated for 2 h at 37°C in a CO_2_ incubator. The CX_3_CR1 antagonist, AA-1-2008, was added (10 nM) to the cells 2 h prior to transmigration assays. To quantify the cellular migration, cells were collected and mixed with a specified amount of Count Bright absolute counting beads (10,000 to 20,000 beads /sample, Molecular Probes). Migration counts were obtained by using a BD LSRII Fortessa cell analyser (BD Bioscience, Franklin Lakes, New Jersey, USA) or AttuneNxT (Thermo Fischer, Waltham, Massachusetts, USA) with FSC/SSC gating on neutrophils/monocyte and beads, respectively, to obtain the total number of cells that had migrated. FlowJo software version 10.5.3 (Tree Star, Ashland, Oregon, USA) was used for flow cytometry analyses.

### Directed migration in µ-slide chemotaxis assay

For the migration chamber experiments, freshly isolated monocytes stained with PKH26 (Sigma-Aldrich) according to manufacturer’s instructions were resuspended in a collagen mixture. Collagen mixture was made by mixing 20µl 10X DMEM (ThermoFisher), 5µl 1M NaOH, 122µl H_2_O, 3µl 7,5% NaHCO_3_, 50µl DMEM and 50µl 3mg/ml Collagen I. This was mixed with 4.5 x 10^5^ monocytes in a 50µl cell suspension, resulting in a final concentration of 1.5 x 10^6^ cells/ml. Then, 6µl of cell-collagen mixture was allowed to polymerize in the µ-slide chambers (ibidi). Before starting the image acquiring process, medium with (+, 100 ng/ml) or without (-) CX_3_CL1 was added to the side-chambers that were in contact with the polymerized cell-collagen mixture. “+/−“ indicates chemokine added to one side-chamber, “+/+” indicates chemokine added in both side-chambers and “−/−“ indicates no chemokines in either side-chamber. One image every 5 minutes was taken, and the µ-slide was kept at 37°C and 5% CO_2_ during the whole imaging process (2 h).

### Flow cytometry analysis

Cells were washed in FACS buffer (2% FCS, 1mM EDTA in PBS) and stained with Zombie UV viability dye and fluorochrome-conjugated mAbs specific (all from Biolegend, San Diego, California, USA) for CD45 (clone HI30, Alexa 700), CD83 (HB15e, Brilliant Violet 421), CD14 (clone M5E2, Brilliant Violet 585), CD56 (HCD56, Brilliant Violet 510), CD19 (HIB19, Brilliant Violet 510), CD3 (OKT3, Brilliant Violet 510), PD-L1 (clone 29E.2A3, Brilliant Violet 605), CD80 (clone 2D10, PE) and HLA-DR (clone Y1/82A, PE/Cy5) for 30 minutes on ice. Cells were then fixed using Cytofix (BD) for 10 minutes and resuspended in FACS buffer. For analysis, monocytes enriched from buffy coats were identified using FCS/SSC and then selected for single live cells positive for CD45 and HLA-DR, negative for CD3, CD56, CD19, and positive for CD14 (Supplementary Fig. 6A). Samples were analyzed with Fortessa LSRII SORP flow cytometer (BD Biosciences), and data were processed with FlowJo 9.7.6 software (Tree Star, Ashland, OR). The geometric MFI of CD83, CD80, CD86 and PD-L1 on the selected monocytes was then reported.

### Phagocytosis and intracellular killing assay

Isolated monocytes were resuspended in complete RPMI media with 2mM L-glutamine and 5% FCS (Gibco) and seeded in a 24-well cell culture plate at 5 x 10^5^ cells/well. Monocytes were then stimulated with rCX_3_CL1-10ng/ml or lung conditioned medium (1:20 dilutions) 16-18 h 37°C in a CO_2_ incubator. For phagocytosis assay, cultured monocytes were infected with *S. aureus* strains (Cowan I and LE2332) at MOI 10 for 60 minutes. The extracellular bacteria were killed by adding flucloxacillin (10mg/ml) and lysostaphin (2ug/ml) and incubated for 20 minutes. The number of phagocytosed bacteria was determined by lysing monocytes in H_2_O followed by serial dilution of lysates, which were then plated on a Columbia blood agar plate incubated at 37 °C for 18–24 h. The bacterial CFU counts were determined the next day. Percentages of phagocytosed bacteria were analyzed and calculated in relation to the inoculum. Similarly, for the intracellular killing assay, stimulated monocytes were infected with *S. aureus* strains at MOI 10 for 60 minutes. After the initial infection of 60, minutes extracellular bacteria were killed by adding flucloxacillin (10mg/ml) and lysostaphin (2ug/ml). The infected monocytes were further incubated for 4 h with the antibiotic- and lysostaphin-containing RPMI media at 37°C in a CO_2_ incubator. Finally, samples from each well were collected, cells were lysed and serially diluted in sterile H_2_O, and then plated on Columbia blood agar plates that were subsequently incubated at 37 °C for 18–24 h. The bacterial CFU were determined the next day. Percentage of intracellular surviving bacteria was analyzed and calculated in relation to the invasion (60 min time point).

### Assays for ROS and NO detection

Monocytes stimulated with rCX_3_CL1-10ng/ml for 18 h at 37°C in a CO_2_ incubator were infected with *S. aureus* strains (Cowan I and LE2332) at MOI 1 for 90 minutes ROS and NO production by monocytes was quantified by using CellROX Green (ThermoFisher) and DAF-FAM Diacetate (ThermoFisher), according to the manufacturer’s protocol. For analysis flow cytometry was used and monocytes were identified as described in Supplementary Fig. 6A. Samples were analyzed with Invitrogen AttuneNxT flow cytometer (ThermoFisher Scientific). The geometric MFI of ROS and NO on selected monocytes is reported.

### Image flow cytometry

For imaging flow cytometry, monocytes were purified and infected with *S. aureus* Cowan producing GFP (MOI 10) for 1 h or 4 h as described above. Cells were fixed in 4% paraformaldehyde and resuspended in 25 μl PBS. Draq5 (Invitrogen) was added at a final concentration of 5 μM, and 10^4^ cells were acquired using an Amnis ImageStreamX Mk II system (Luminex, Austin, Texas, USA). Using the IDEAS 6.0 software (Luminex), the number of bacteria per cell was analysed essentially as previously described using the feature Spot count_Peak [(M02, Staph, Bright, 6) (58)]. The mean spot count in the monocyte population, reflecting the average number of bacteria per cell, is reported. The intracellular killing of GFP-producing Cowan I was estimated by calculating the percentage of bacteria that were detectable 4 h post infection.

### Statistical analysis

Data were analyzed with GraphPad Prism software v.8.1.1 (GraphPad Software Inc, San Diego, California, USA) and the statistical methods used are indicated in the respective figure legend. Differences were considered to be statistically significant at *p* <0.05. For all statistical analysis *p<0.05, **p<0.01 and ***p<0.001.

### Data availability

The authors declare that the data supporting the findings of this study are available within the article and its Supplementary Information files, or from the corresponding author on request.

## ACKNOWLEDGEMENTS

The authors thank all patients, and volunteers for participating in this study and, all clinical staff involved in collection of blood and airway samples. Our work was supported by grants from the Karolinska Institutet and Stockholm County council, as well as the Swedish Governmental Agency for Innovation Systems (VINNOVA) under the frame of NordForsk (Project no. 90456, PerAID), the Swedish Research Council under the frame of ERA PerMed (Project 2018-151, PerMIT), and by a project grant from the Swedish Research Council (521-2014-6722). PC received an MD-PhD scholarship from the Karolinska Institutet. ML was supported by the Swedish Children’s Cancer Foundation. S.M.S was supported by a grant from the Swedish Society for Medical Research (SSMF, P17-0179). S.M.S and A.S.Z was supported by the grant from Uniscientia Foundation grant. A.S.Z was supported by the Swiss National Science Foundation (31003A_176252) and the Clinical Research Priority Program (CRPP) of the University of Zurich, Switzerland. A.W. was supported by a grant from the Swedish Research Council (2014-396). Imaging and flow cytometry at University of Zurich were performed using equipment of the Centre for Microscopy and Image Analysis as well as the Flow Cytometry Facility. We would like to thank Dr. Gayathri Arakere, Society for Innovation and Development, Indian Institute of Science, Bengaluru, India for generously providing us with the lung Infection isolates (NP796 and LE2332) and the reference CA-MRSA strain (USA300). The authors declare no conflict of interest, financial or otherwise.

## AUTHOR CONTRIBUTIONS

SMS, PC and RMB were responsible for and performed the *in vitro* experiments and sample analyses; SC, KS, RAS and ASZ, cared for patients and provided clinical data; SC, VÖ, KS, RAS and ASZ provided patient samples; EMM contributed with stained and scanned lung tissue samples; RMB contributed to monocyte migration assays and sample analyses; AW contributed with imaging flow cytometry; MH contributed to the phagocytosis, transwell migration, hemolysis assays, enzymatic activity assay and in vivo murine infection experiments; AGM and TAS contributed to the mice approvals and performing in vivo murine infection experiments and CX3CL1 ELISA; JS contributed to fluorescence microscopy analyses; SBB and ML contributed to monocyte stimulation assays and ELISA; TSC and VT performed the *in vivo* experimental models and associated analyses; SMS, PC, ANT, and MS initiated the study, and contributed to the study design and analysis and interpretation of the results; MS supervised the study; PC and SMS prepared the figures; SMS, PC, and MS wrote the manuscript and all authors participated in data interpretation and editing and finalizing the manuscript, which was approved by all authors.

