## Supplementary Figures for "Alpha-toxin elicited CX_3_CL1-release via ADAM10 in *Staphylococcus aureus* pneumonia impairs bactericidal function of human monocytes"

**This PDF file includes:**

Supplementary **Figure 1**

Supplementary **Figure 2**

Supplementary **Figure 3**

Supplementary **Figure 4**

Supplementary **Figure 5**

Supplementary **Figure 6**

Supplementary **Figure 7**

Supplementary **Figure 8**

Supplementary **Table 1**.

Supplementary **Table 2**.

Legend for Supplementary **Video.**

**Other supplementary materials for this manuscript include the following:**

Supplementary Video

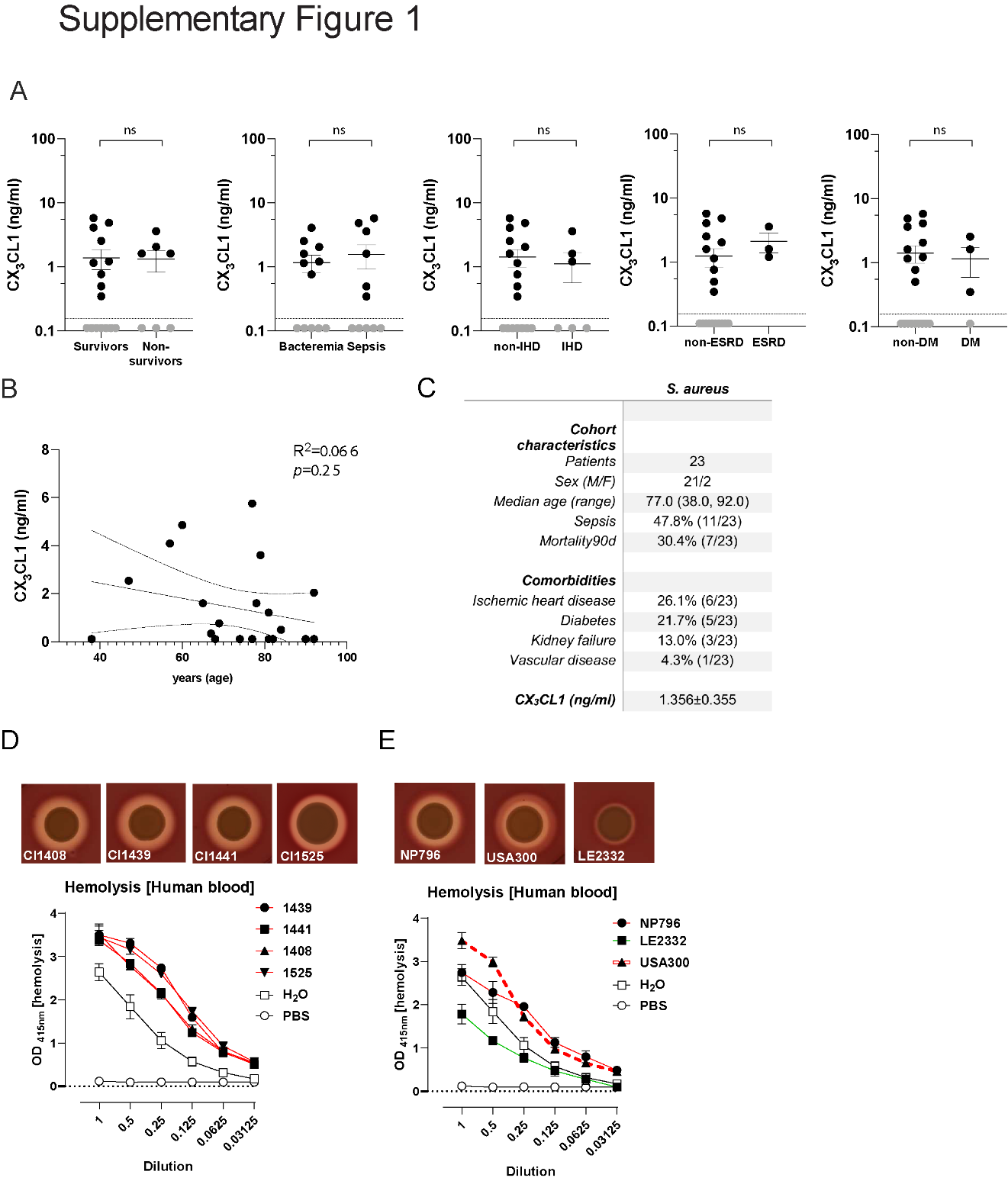

Supplementary Fig. 1. CX3CL1 plasma levels of bacteremia cohort and blood hemolysis assays. A) *S. aureus*-positive patients were divided into groups based on survivors and non-survivors, sepsis and bloodstream infection without sepsis (bacteremia), ischemic heart disease (IHC), end-stage renal disease (ESRD) and diabetes mellitus (DM). The levels of CX_3_CL1 were determined by ELISA with at LOD of 0.156 ng/ml. Statistical significances were determined by Mann-Whitney Unpaired t-test. B) Linear regression analysis of age and CX_3_CL1 levels. C) Characteristics of bacteremia cohort. Demographic data and comorbidities of patients with *S. aureus* infection. The level of CX_3_CL1 was determined by ELISA and the mean value ± SEM is shown. Source of infection included arthritis/osteomyelitis (n=8), blood stream infection (n=3), endocarditis (n=5), skin/soft tissue infection (n=2), stentgraft infection (n=1) and unknown (n=4). D) Images show the zone of hemolysis in blood agar, and diagrams show the percent lysis of human red blood cells by clinical *S. aureus* strains (NP796 and LE2332) and USA300 strains. E) Representative images show representative pictures of the zone of hemolysis in blood agar, and diagrams show the percent lysis of human red blood cells by four clinical *S. aureus* strains from patients with *S. aureus* respiratory tract infection (Fig 1A). Red blood cells lysis was evaluated by determining the absorbance at 415nm. PBS alone and H_2_O were used as negative and positive controls, respectively.

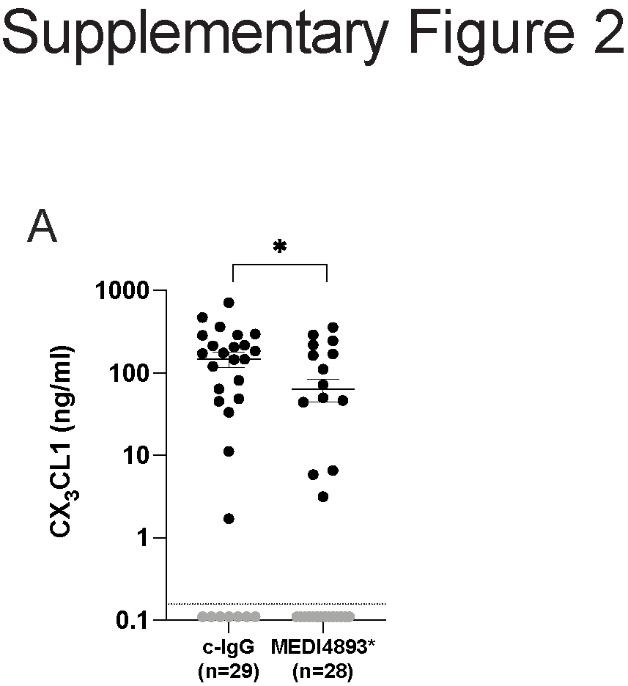

Supplementary Fig. 2. *In vivo* blockade of α-toxin using an anti-α-toxin antibody. CX_3_CL1 in bronchoalveolar lavage of S. aureus infected mice treated with the anti-α-toxin antibody MEDI4893* or an isotype matched control-IgG-antibody. Mice were infected with the *S. aureus* strain USA300, dosed at (5 x 10^7^ CFU) intranasally and bacterial suspensions were administered in 50 µL of PBS. MEDI4893* or c-IgG was administered in 0.5 mL intraperitoneally (IP) 24h prior to infection. Animals were euthanized with CO_2_ 24h post infection, and BAL fluid was collected for CX_3_CL1 measurement. The levels of CX_3_CL1 were determined by ELISA. Values below LOD were set to 0.1103 ((LOD)⁄(√2)) and are shown in grey. Statistical significances were determined by Mann-Whitney Unpaired t-test.

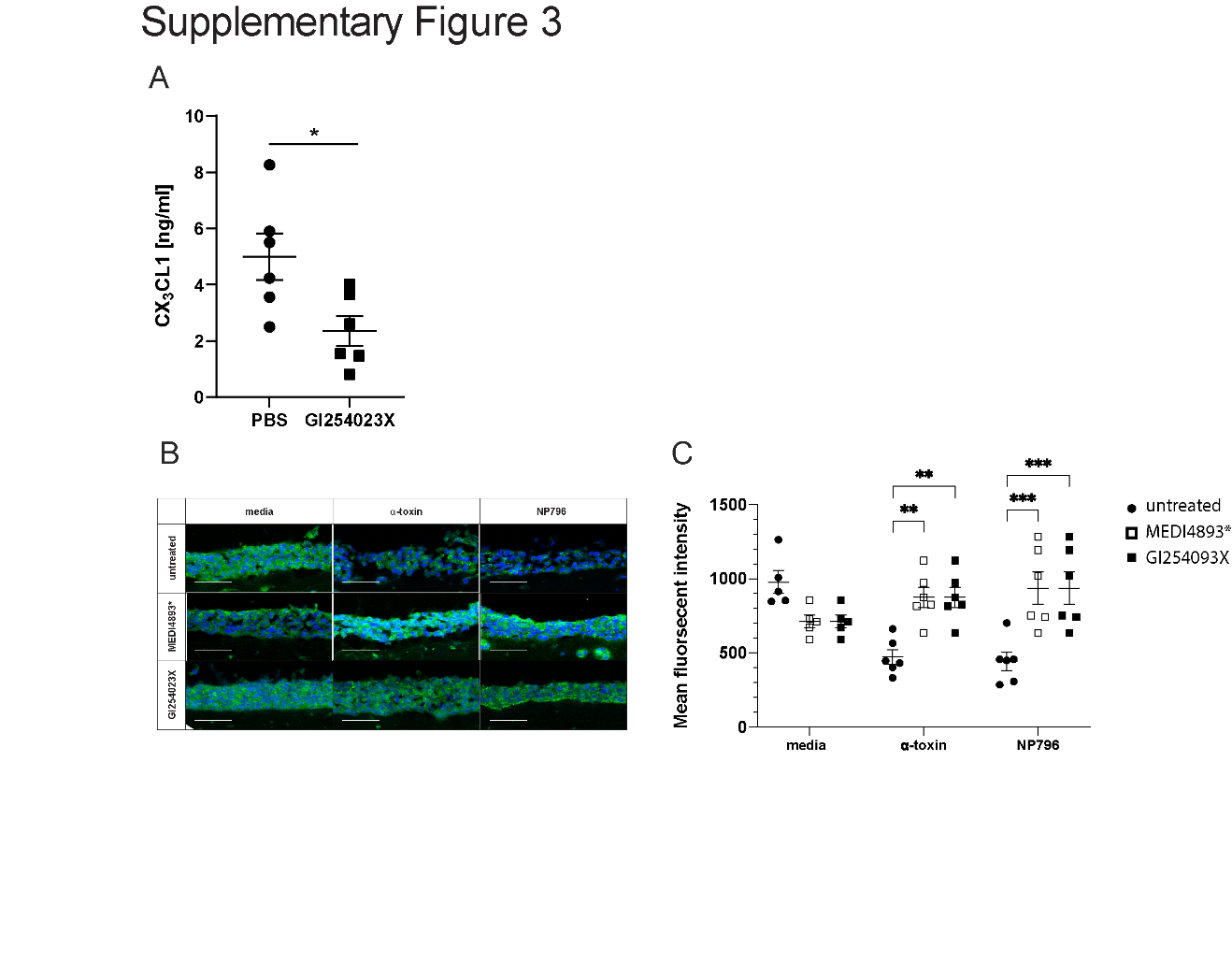

**Supplementary Fig. 3. In vivo blockade of α-toxin-mediated effects and E-cadherin in lung model stimulated with α-toxin and NP796 bacterial culture supernatant**. A) CX_3_CL1 in bronchoalveolar lavage of mice infected with the *S. aureus* strain USA300, dosed at (5 x 10^7^ CFU) and pre-treated with the ADAM10 inhibitor, GI254023X at 200mg/kg administered in 100µl volume intraperitoneally (IP) for every day 3 to 4 days prior infection. PBS treatment was used as control. Mice were infected intranasally and bacterial suspensions were administered in 25 µL of PBS. Animals were euthanized with CO_2_ 24h post infection, and BAL fluid was collected for CX_3_CL1 measurement. The levels of CX3CL1 were determined by ELISA. Statistical significances were determined by Mann-Whitney Unpaired t-test. B) Immunofluorescence staining of E-cadherin (green) and cell nuclei (blue, DAPI) in sectioned lung model stimulated with *S. aureus* α-toxin (100 ng/ml) or NP796 bacterial culture supernatants (1:100), in the absence (untreated) or presence of anti-α-toxin antibodies (200 µg/ml, MEDI4893*), or the ADAM10 inhibitor, GI254023X (10 µM). C) The mean fluorescent intensity in the epithelial area (as indicated by DAPI) in sectioned lung model, stimulated with *S. aureus* α-toxin (100 ng/ml) or NP796 bacterial culture supernatants (1:100), in the absence (untreated, an isotype matched antibody or DMSO was used, filled circles) or presence of anti-α-toxin antibodies (200 μg/ml, MEDI4893*, open squares), or the ADAM10 inhibitor, GI254023X (10 µM, filled squares). Images from a minimum of five independent experiments were analysed. Statistical significances were calculated by ordinary two-way ANOVA, with uncorrected Fisher’s LSD.

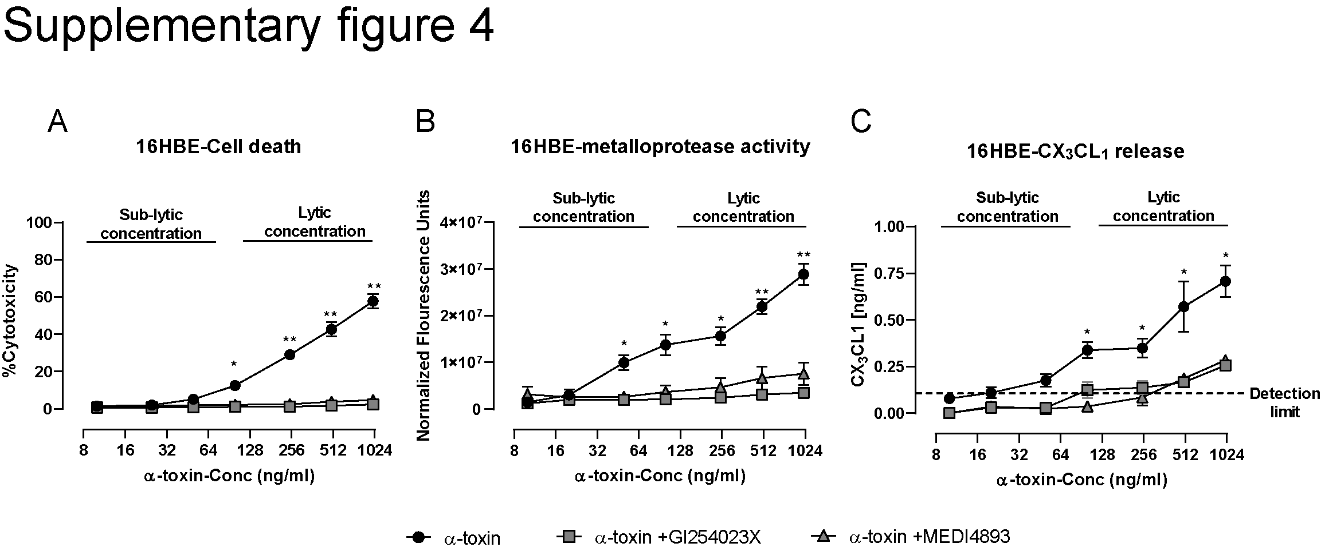

**Supplementary Fig. 4 Dose dependent effect of alpha toxin-induced CX_3_CL1 release.** A-C) Lung epithelial cells (16HBE) were exposed to 10, 25, 50, 100, 250, 500 1000 ng/ml of α-toxin in the absence (filled circles) or presence of anti-α-toxin antibodies (200 μg/ml, MEDI4893*) or the ADAM10 inhibitor, GI254023X (10 µM). 16HBE cells were pre-treated with GI254023X for 4h or α-toxin was preincubated with MEDI4893* for 4h before stimulations. A) Cytotoxicity was determined by LDH measurements assessed in supernatants of lung epithelial cells (16HBE). 16HBE cells were exposed for 2h after which OD values were obtained. Percent cytotoxicity as related to the positive control (Triton lysed cells) is shown. B) Metalloprotease enzymatic activity was determined on the basis of cleavage of Mca-Pro-Leu-Ala-GlnAla-Val-Dpa-Arg-Ser-Ser-Ser-Arg-NH2 fluorogenic peptide substrate. The data is presented as normalized fluorescence value to that of unstimulated cells C) ELISA determination of CX_3_CL1 levels in culture supernatants of 16HBE exposed to α-toxin for 2 h. All experiments were performed at least three times. In A to C, data are represented as the mean value ± SEM. The statistical significances were calculated by Kruskal-Wallis test with Dunn’s multiple comparison

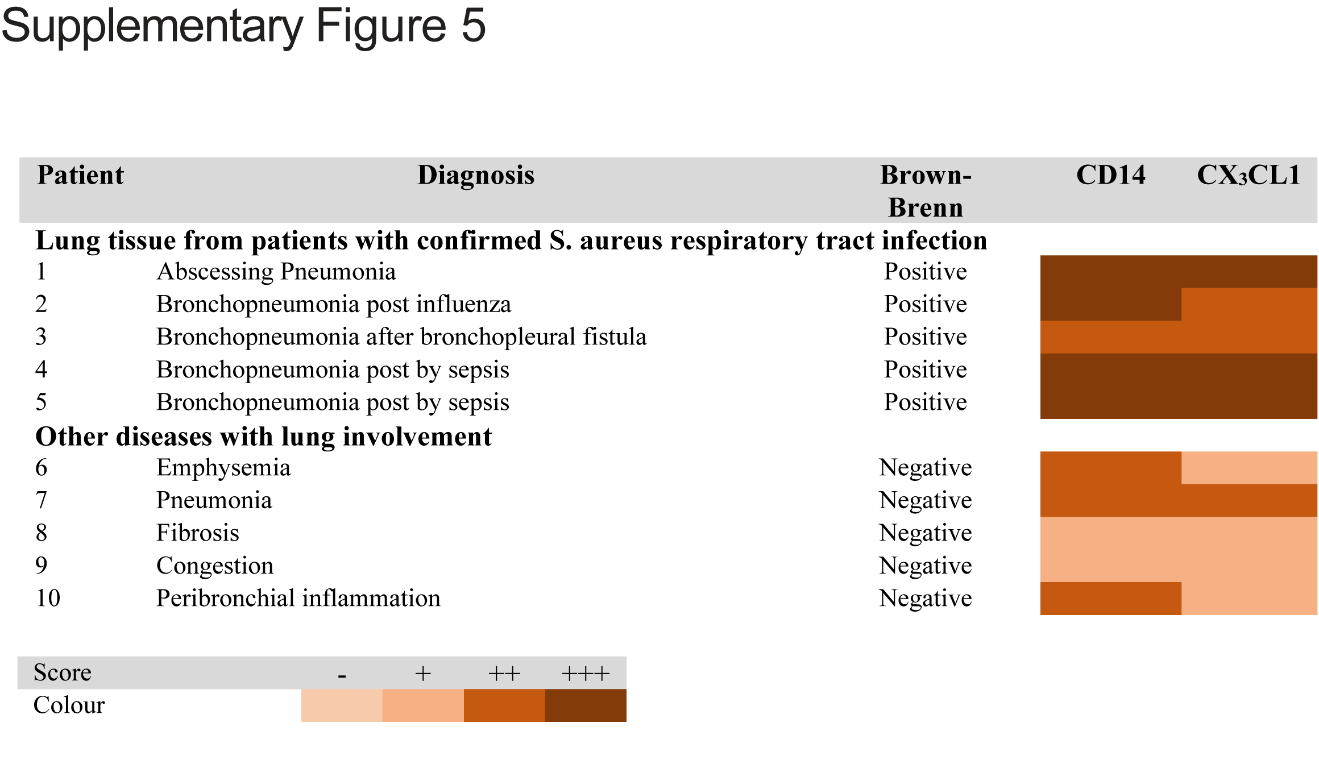

**Supplementary Fig. 5. Summary of histological and immunochemical qualitative analysis of lung tissue biopsies.** Lung tissue biopsies from patients with *S. aureus* respiratory tract infection (n=5) or control tissue samples from patients with lung diseases other than infection (n=5) were stained with Brown-Brenn reagents or for CD14 and CX_3_CL1. A summary of the Brown-Brenn and **immunohistochemical characteristics of the stained lung tissue samples is shown.** For the Brown-Brenn staining, biopsies were scored positive or negative. For the CD14 and CX_3_CL1 staining, biopsies were scored based on immunostaining intensity: - (negative), + (low), ++ (intermediate), or +++ (high). A summary of the scoring in relation to diagnosis is shown.

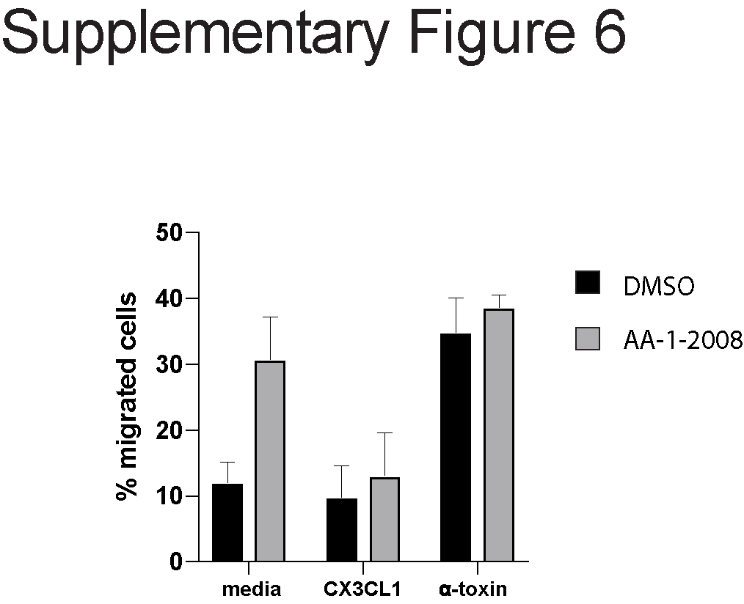

**Supplementary Fig. 6. Neutrophil migration in transwells**. Human neutrophils were isolated from fresh whole blood obtained from healthy volunteers using Polymorphprep (Axis-Shield, Oslo, Norway) by performing density gradient separation according to the manufacturers’ instructions. Migration of neutrophils for 2 h in a transwell assay in response to supernatants from unstimulated lung model (media), lung models stimulated for 24 h with α-toxin (50 ng/ml), or medium supplemented with recombinant CX_3_CL1 (10 ng/ml), in the absence (DMSO, diluent of AA-1-2008, black bars) or presence of the CX_3_CL1 receptor antagonist (AA-1-2008, 10 nM, grey bars) was assessed. Pre-treatment with the CX_3_CR1 antagonist AA-1-2008 (10nM, grey) or DMSO (vehicle, black) was for 2 h prior to the migration assay. Statistical significances were calculated by ordinary two-way ANOVA, with Bonferroni’s multiple comparisons test.

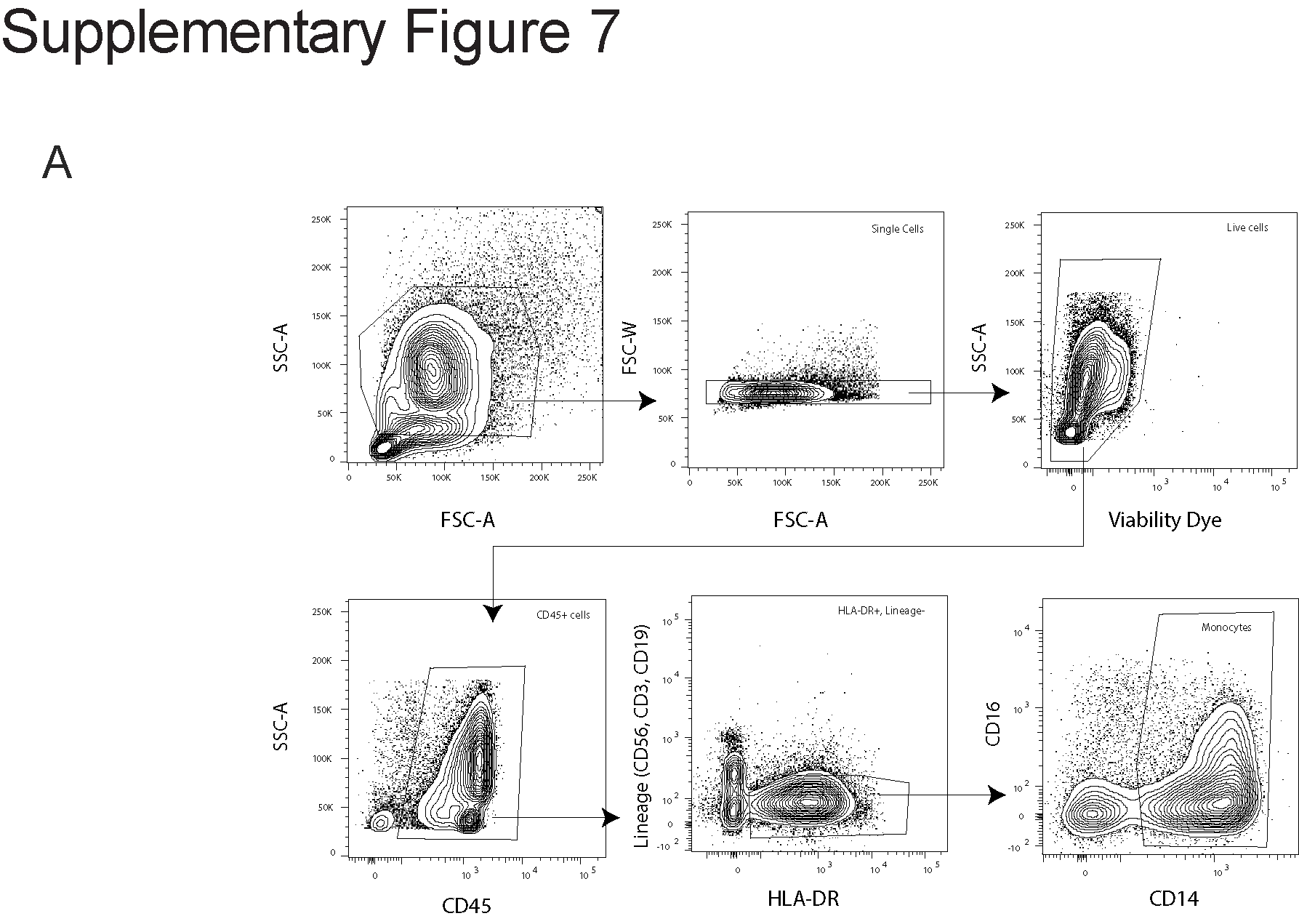

**Supplementary Fig. 7. Monocyte gating strategy.** A) Mononuclear leukocytes were identified in monocyte enriched fractions using FCS/SSC. Then single (FSC-W/FSC-A) live cells (SSC-A/Viability dye) were further gated to select CD45 positive cells negative for CD3, CD56, and CD19, while positive for HLA-DR and CD14 with a heterogenous expression of CD16.

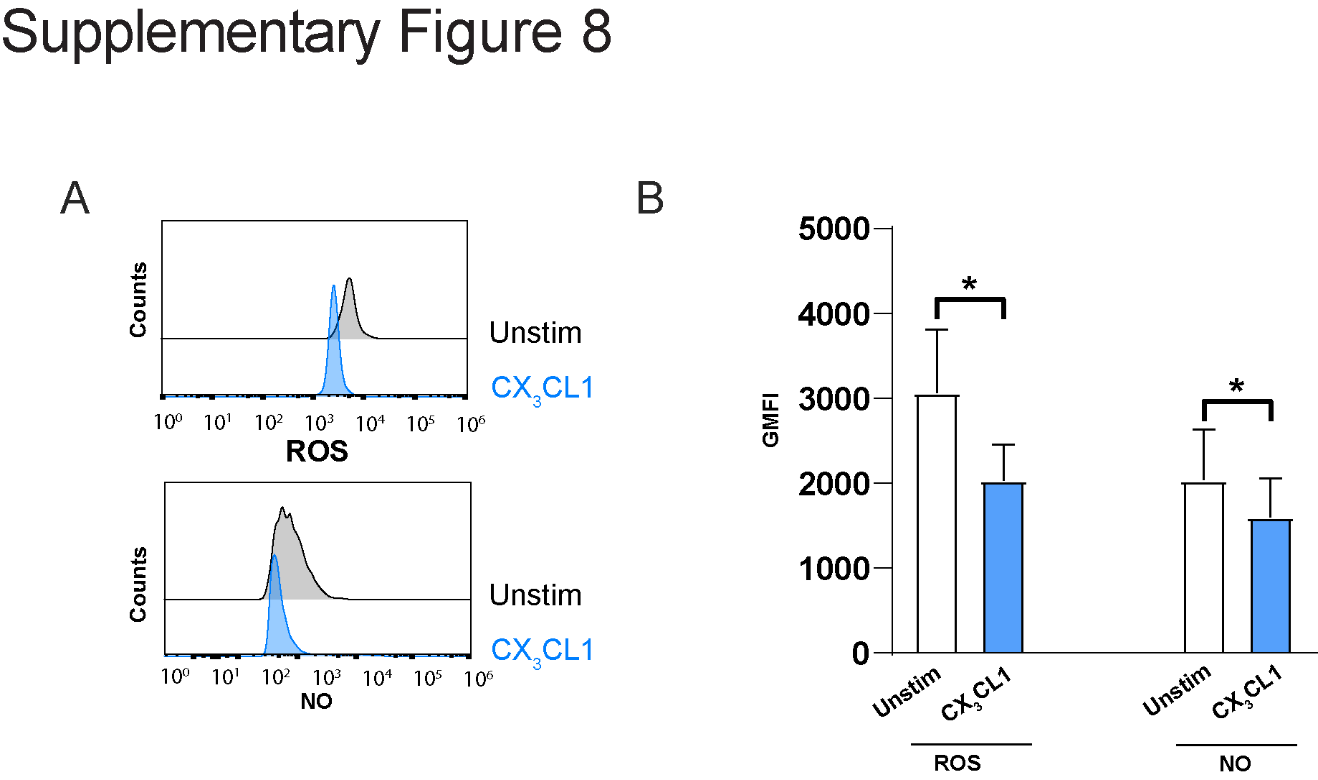

**Supplementary Fig. 8. Monocyte respiratory burst.** A-B) Monocytes stimulated with CX_3_CL1 (10 ng/ml) for 18 h or left unstimulated, were infected with *S. aureus* strain Cowan I (MOI 1), and analysed for ROS and NO production. Representative histograms (A) of ROS and NO production, and representative bar graphs (F) of ROS and NO production by Cowan I-infected monocytes. The graphs show the mean ± SEM of five independent donors. In B statistical significances were calculated using Wilcoxon matched-pair signed-rank test.

Supplementary Table 1. Characteristics of the patients included in the study divided based on the methods performed and the category of infection.

| **Method** | **Patients and** | | **CTRL** | ***S. aureus*** |
| --- | --- | --- | --- | --- |
| **CX_3_CL1 ELISA**  respiratory tract infection | | **samples analyzed**  Number of patients | 4 | 4 |
|  | | BAL | 1 | 2 |
|  | | Tracheal aspirates | 3 | 2 |
|  | | Age* | 44.25 ± 16.03 | 42.00 ± 12.39 |
|  | | Sex (F/M) | 1/3 | 0/4 |
| respiratory tract infection | | Number of patients | 8 | 4 |
|  | | Plasma | 8 | 4 |
|  | | Age* | 33.38 ± 4.33 | 42.00 ± 12.39 |
|  | | Sex (F/M) | 4/4 | 0/4 |
| bloodstream infection | | Number of patients | 19 | 23 |
|  | | Plasma | 19 | 23 |
|  | | Age* | 46.78 ± 12.15 | 74.09 ± 13.89 |
|  | | Sex (F/M) | 8/11 | 2/21 |
| **Histology/Histochemistry** | |  |  |  |
| respiratory tract infection | | Number of patients Lung (Resection) | 5  3 | 5  3 |
|  | | Lung (Autopsy) | 0 | 2 |
|  | | Age* | 72.66 ± 11.46 | 59.60 ± 13.06 |
|  | | Sex (F/M) | 1/2 | 4/1 |

CTRL = controls without *S. aureus* infection and healthy donors; F = female; M = male. *Mean ± standard deviation.

| **Patient Number** | **Clinical Diagnosis/** | | **Bacteremia** | | **Mortality** |
| --- | --- | --- | --- | --- | --- |
| **ICU Control group** | | **Presentation** |  | |  |
| 1 | | Cardiogenic shock | | No | Yes |
| 2 | | Peritoneal carcinomatosis with ventilator pneumonia | | No | Yes |
| 3 | | Primary acute respiratory distress syndrome | | No | Yes |
| 4 | | Bilateral lung transplantation | | No | Yes |
| ***S. aureus* respiratory tract infection** | |  | |  |  |
| 1 | | Hyper-IgE syndrome with acute lung infection causing pneumonia | | No | No |
| 2 | | Right sided endocarditis with pulmonary emboli and abscess | | Yes | Yes |
| 3 | | Septic shock and bilateral pneumonia with influenza A | | No | No |
| 4 | | Right sided endocarditis with sceptic emboli to the lung | | Yes | No |

Supplementary Table 2. Clinical diagnosis and presentation at time of sampling of the patients included in the study

Legend for Supplementary Video

Movie S1 (separate file). Directed migration of monocytes. Monocytes were allowed to migrate in a polymerized collagen mixture. The chamber holding the cell-collagen mixture is connected to side-chambers holding DMEM with (+, 100 ng/ml) or without (-) CX_3_CL1. An automated cell-tracking algorithm is overlaid on the video to represent the distance and direction travelled for each monocyte.
